# High-throughput 3D super-resolution ultrasound imaging

**DOI:** 10.1101/2025.08.20.671215

**Authors:** Weisong Zhao, Nanchao Wang, Zhenqian Han, Xining Xu, Xiangyu Ma, Tianhua Zhou, Jiahui Gui, Yuzhen Liu, Qianbo Liu, Liying Qu, Wenhao Liu, Xiangyan Ding, Xin Liu, Dean Ta, Jiubin Tan, Liangyi Chen, Haoyu Li

**Author notes:** These authors contributed equally to this work (Weisong Zhao, Nanchao Wang, Zhenqian Han).

## Abstract

Capturing fast hemodynamics in deep organs is essential for understanding microvascular regulation of organ responses to physiological demands and pathological stress. However, non-invasive three-dimensional (3D) imaging of these microscale processes remains challenging due to trade-offs between spatial resolution, imaging speed, and penetration depth. Here, we present fluctuation-based high-order super-resolution acoustic microscope (FLAME), a tracking-free 3D ultrasound imaging technology capable of fast microvascular angiography and flow measurement. Using as few as 30 volumes, FLAME improves 3D ultrasound resolution by 8-fold (∼50 μm) and shortens data acquisition time by over two orders of magnitude, from tens of seconds to tens of milliseconds, compared with conventional tracking-based super-resolution approaches^1-6^. This high-throughput capability supports high-fidelity imaging of tissue functions under flexible experimental settings. FLAME achieves an unprecedented 3D super-resolution frame rate of ∼40 Hz, capturing transient hemodynamic responses to various physiological and pathological stimuli in different mouse organs, from the microvascular level to the whole-body scale. With open-sourced implementation, FLAME provides an accessible platform for real-time, in-depth hemodynamic monitoring across diverse biomedical applications and clinical translation.

## INTRODUCTION

Optical super-resolution microscopy offers unprecedented spatial resolvability^7-11^, but the reliance on light excitation and modulation restricts its applicability to thin, transparent samples. In opaque tissue, strong photon scattering severely constrains imaging depth and field of view (FOV), making large-scale, high-throughput imaging challenging. Unlike light, tissues allow ultrasound (US) to pass through more clearly. By further introducing US contrast agents—intravenously injected microbubbles (MBs)—contrast-enhanced US (CEUS) imaging^12^ can highlight the overall vascular network^13,14^. However, governed by wave diffraction, its spatial resolution typically remains at the scale of the acoustic wavelength, which is insufficient to resolve the detailed microvascular structures and hemodynamics.

Ultrasound localization microscopy (ULM)^1,2,4^ adapts the principles of stochastic super-resolution optical microscopy^7,8^ by localizing and tracking the flowing MBs to surpass the acoustic diffraction limit by up to ∼10-fold, visualizing the microvascular networks in detail. However, its tracking mechanism requires precise localization of sparsely distributed MBs in each image frame, inherently enforcing a long data acquisition time on the level of minutes^2,3^ (**Fig. 1a**). Practically, ULM applications face several critical challenges^15^: (i) The slow accumulation of MB localizations requires tens of thousands of frames, yielding essentially time-averaged hemodynamic maps; (ii) For transcranial or deep-tissue imaging, acoustic aberrations distort the acoustic signals, impairing MB isolation and tracking precision; (iii) For quantitative imaging of the blood flow speed, there is an inherent trade-off between the accumulation of slowly-moving MBs and the accurate detection of fast-moving MBs, resulting in an overall imaging fidelity degradation; (iv) For clinical translation, the single bolus injection of MBs with high concentrations often leads to the spatial overlapping of MBs, resulting in deteriorated localization accuracy. To sum up, these technical challenges are particularly critical for 3D imaging applications in hemodynamic studies that demand high temporal resolution on the sub-second scale^5,16^.

**Fig. 1.**
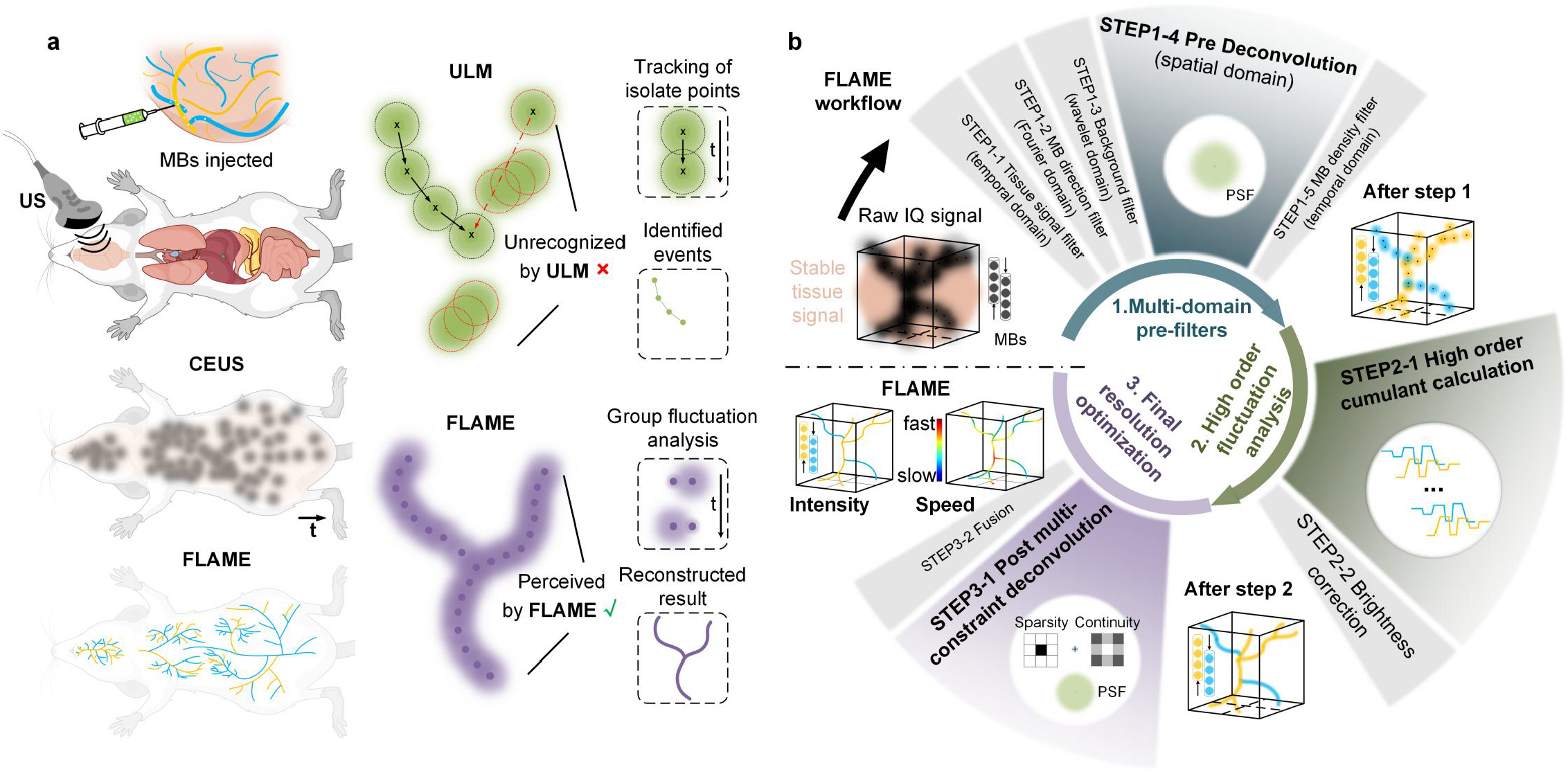
Principle of FLAME. **a**, Schematics of the experimental setup for CEUS and FLAME imaging (left) and conceptual differences between ULM and FLAME (right). After intravenous injection of MBs, a volumetric transducer collected US scattering signals from both stationary tissues and flowing MBs. By executing an SVD filter over time, CEUS imaging results (power Doppler images) were obtained, containing isolated MB signals (black) and residual tissue components (flesh color). Finally, beyond the SVD filter, FLAME reconstruction was applied to resolve microvasculature at a super-resolution scale (∼8-fold finer). In contrast to ULM requiring localizing and tracking individual MB flowing events, FLAME leverages group-level fluctuation analysis to generate integrated reconstruction results. Unlike ULM, which relies on precise localization and tracking of individual MB flow events, FLAME harnesses group-level fluctuation analysis to directly yield super-resolution vascular reconstructions. **b**, Reconstruction workflow of FLAME (detailed version in Extended Data Fig. 1). The input raw time-lapse IQ volumes underwent three major processing steps to generate super-resolution volumes of both intensity and flow speed. [Schematics were created with www.figdraw.com]

To moderate these challenges, fluctuation-based super-resolution US imaging methods have been reported^17-22^. Similar to super-resolution optical imaging strategies^9,23,24^, the flow of MBs along the vessels results in signal intensity fluctuations of the CEUS images. Instead of localizing the MBs as in ULM, the fluctuation-based methods analyze the temporal statistics of the high-density MB movements, which can effectively shrink the point spread function (PSF) width. However, these methods usually need hundreds to thousands of CEUS frames and are excessively sensitive to noise levels. Moreover, the limited usage of high-order statistics restricts the improvement in three-dimensional (3D) resolution. To date, non-invasive high-throughput imaging of microscopic hemodynamics in deep tissues remains an open quest.

Here, we present FLAME (FLuctuation-based high-order super-resolution Acoustic MicroscopE), a volumetric super-resolution technique for CEUS imaging. Our recent development of an ultrafast super-resolution optical method made this technologically possible^11^. FLAME simultaneously maximizes the volumetric imaging speed (using 30 volumes) and the resolution enhancement (up to 8-fold) in all three spatial dimensions. FLAME allows for high-fidelity vascular mapping with two orders of magnitude less data acquisition time than classical tracking-based methods, while delivering comparable resolution enhancement. Implemented on a 4-MHz hemispherical US array, FLAME achieved an FOV of approximately 1 cm^3^ with ∼50 μm resolution and ∼40 Hz volumetric frame rate, providing similar speed with cutting-edge 3D functional US imaging^25-27^ but superior spatial details. We thoroughly characterized FLAME on *in vitro* flow phantoms with different experimental conditions, validating its high accuracy and effectiveness. To illustrate its versatility in biomedical applications, we applied FLAME to various traditionally challenging tasks in small animal models, including (i) transcranial whole-brain imaging under low signal-to-noise ratio (SNR); (ii) transient hemodynamic imaging in response to pharmaceutical and neuromodulatory stimuli; and (iii) high-throughput whole-body and deep-organ imaging.

## RESULTS

### Principle of FLAME

Unlike ULM, FLAME leverages the assembled temporal blinking/fluctuations of group MBs rather than tracking the locations of single ones (**Fig. 1a**). It progressively enhances spatial resolution and image quality through a series of purpose-specific volumetric processing steps, each tailored to maximize its capability. By combining multi-level filtering, constrained deconvolution, and brightness correction, FLAME enables high-order stochastic calculations to achieve 3D resolution enhancement from a small number of image frames. We provide an integrated and versatile solution consisting of three major steps as detailed below (**Methods, Fig. 1b, Extended Data Fig. 1, Supplementary Video 1**).

First, to extract assembled motion events of group-distributed MBs from the noisy US volume sequences, we designed a series of multi-domain pre-filters to maximize the effective fluctuation contrast ratio. The raw IQ (in-phase/quadrature-phase) data containing MB motion signatures undergo cascaded filtering. In STEP 1.1∼1.3, temporal singular value decomposition (SVD)^28,29^ removes stationary tissue signals; Fourier-based directional filtering^30^ separates upward and downward flowing MBs; and wavelet decomposition eliminates residual motion artifacts from large-scale tissue structures. In STEP 1.4, to further shrink the volumetric point spread function (PSF) and suppress out-of-focus signals and noise, we apply a 3D pre-deconvolution^11^ to the filtered images. This core step effectively attenuates random pixel-level fluctuations caused by noise and highlights the temporally correlated fluctuations, hence improving the efficiency of high-order statistical analysis among vessels while preserving frame-to-frame linearity. In STEP 1.5, we adapt the HAWK (Haar wavelet kernel) analysis as a density filter to further increase the temporal separability of MB groups. Originally developed for single-molecule localization microscopy (SMLM)^31^, HAWK can separate emitter populations based on their distinct emission profiles through multi-level temporal bandpass filtering. SMLM typically achieves high emitter density by reducing excitation intensity, which inherently lowers the signal-to-noise ratio (SNR). However, HAWK analysis demands a high SNR to ensure reliable separation, creating a fundamental trade-off in SMLM applications. In contrast, HAWK is particularly well-suited for CEUS, where our preprocessing pipeline consistently provides high-quality input data for downstream statistical analysis. Thus, HAWK in US imaging effectively reduces the MB density per frame, virtually stretching spatial information over time and converting densely overlapping events into temporally resolvable signals within a ∼3× extended image sequence.

Our recent work^11^ identified three fundamental challenges when applying high-order statistical analysis to fluorescence imaging, including 1) the large dynamic range in signal intensity due to fluctuation heterogeneity; 2) the requirement for long temporal sequence; and 3) the requirement for high blinking contrast. In the context of US imaging, we tackle these challenges through a comprehensive solution. In STEP 1, adaptive filtering isolates MB fluctuations with refined spatial resolution, improved SNR, and reduced MB density. This enables high-order statistical analysis in STEP 2.1 to focus on high-contrast temporal fluctuations, allowing reliable 6^th^-order cumulant computation from only 30 US volumes (virtually HAWK-generated 90 volumes). The resultant high-order cumulant images inherently exhibit a large intensity dynamic range, so we perform local dynamic range compression (LDRC)^32^ to preserve microvessel structure integrity and reduce nonlinearity. After that, we maximize the image resolution and fidelity by multi-constraint deconvolution, which incorporates the 3D continuity and sparsity constraints^10^. Finally, we execute a data fusion step to finalize the volumetric reconstructions. The resultant high spatial resolution also enables relative flow speed quantification via temporal standard deviation analysis^33^ of MB fluctuations. Each FLAME volume was reconstructed from 30 US volumes, followed by a 6^th^ root transformation to normalize the intensity distributions. Then, every four FLAME volumes were used to compute one blood perfusion intensity map (by rolling average) and one flow-speed map (by rolling standard deviation), providing high-quality reconstruction with high temporal resolution.

### FLAME unlocks faster and simpler microangiography

We evaluated the performance of FLAME for whole-brain microvasculature in mice with intact skin and skull using a 4-MHz hemispherical array (**Extended Data Fig. 2a**), aiming to explore transcranial deep-brain imaging under low SNR conditions. For benchmarking, we tested two different MB concentrations: a ULM-compatible low-density condition (‘low-density’) for cross-validation against ULM as the reference (**Fig. 2a-2e, Extended Data Fig. 3a**), and a ULM-incompatible high-density condition (‘high density’) for more challenging experimental conditions (**Extended Data Fig. 5**). Unlike the regions of interest (ROIs) in standard CEUS images, FLAME reconstructions by 120 volumes over ∼0.56 seconds (FLAME-120v, **Fig. 2a-2c, Supplementary Video 2**) yielded high-quality super-resolved vessel maps that are comparable to the ULM reconstructions by 4,800 volumes over ∼22.33 seconds (ULM-4,800v, **Fig. 2c**). This result demonstrates FLAME’s superior performance over CEUS and ULM, especially in its simultaneous high spatiotemporal resolutions. Moreover, the spatial resolution achieved by different imaging methods was quantified by the Fourier shell correlation (FSC) analysis^34,35^ (**Fig. 2d**), showing 74 μm for FLAME-30v, 58 μm for FLAME-120v, and 57 μm for ULM-4,800v. In contrast, CEUS had a diffraction-limited resolution of 452 μm, only resolving large vessels but not microvascular details (**Fig. 2a-2d**).

**Fig. 2.**
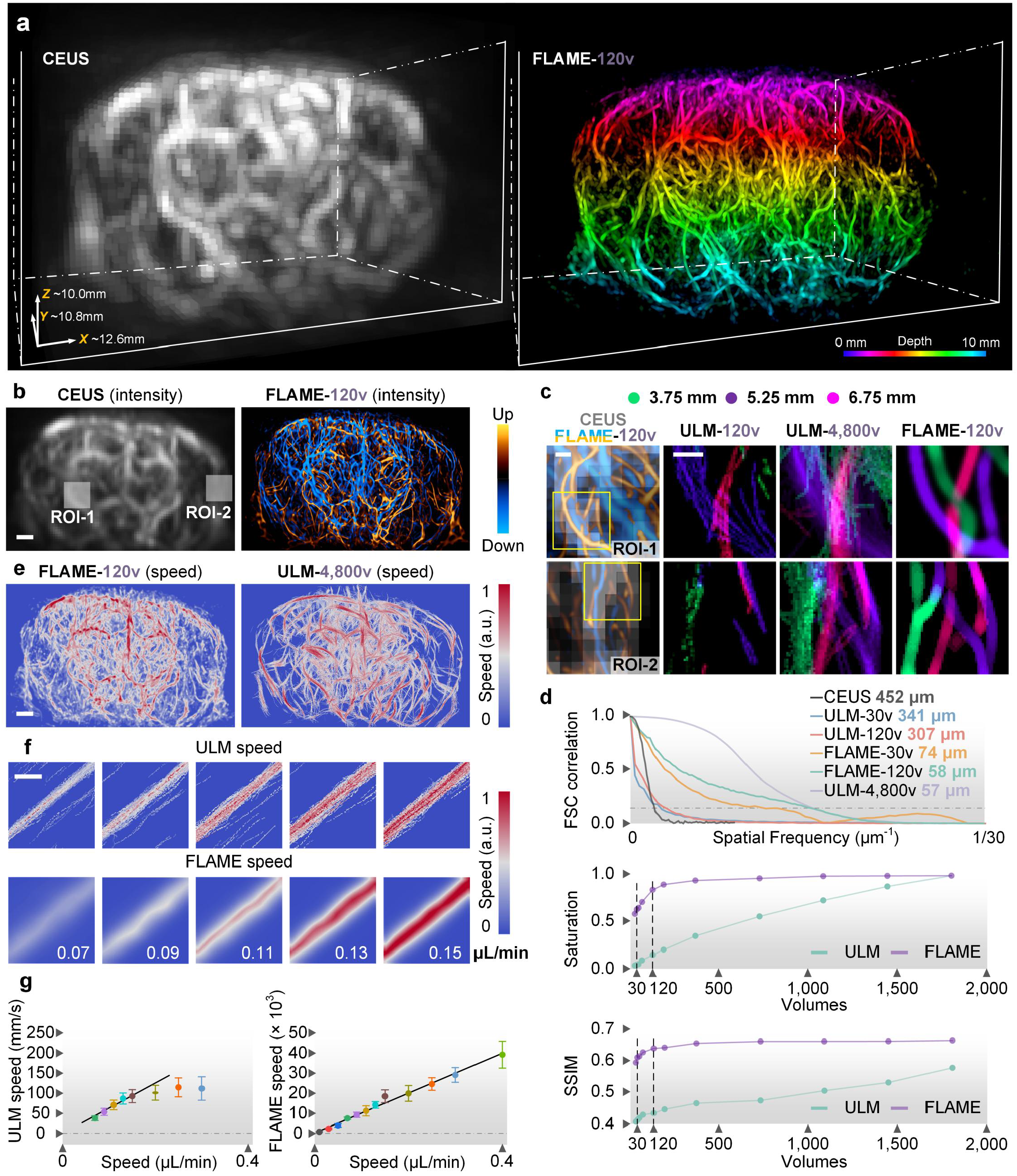
Systematic evaluation of FLAME. **a**, 3D rendering of a representative whole-brain microvascular network in a living mouse with intact scalp and skull recorded by CEUS (left) and 120-volume reconstructed FLAME (FLAME-120v, right). Volume size is labeled at the bottom left corner. **b**, Color-coded 3D distributions of the microvasculature in (**a**) under CEUS (power Doppler image, left) and FLAME-120v (right). Scale bars, 1 mm. **c**, From left to right are magnified views of the white boxes in (**b**) reconstructed by CEUS (gray) merged with dual-channel FLAME-120v (orange and blue) and enlarged regions enclosed by the yellow boxes under ULM-120v, ULM-4,800v, and FLAME-120v. Different colors indicate different depth positions. Scale bars, 0.2 mm. **d**, From top to bottom are FSC resolution analysis, saturation rates, and SSIM values of ULM (turquoise) and FLAME (violet) results reconstructed by different volumes, respectively. SSIM values were estimated from reconstructions against ULM-4,800v. Saturation rates were calculated from randomly selected regions in ULM and FLAME reconstructions against their 1,800-volume results, respectively. **e**, Maximum flow speed projection maps of FLAME-120v (left) and ULM-4,800v (right). Scale bars, 1 mm. f, Flow speed maps measured by ULM-3,000v (top) and FLAME-120v (bottom) using a tube *ex vivo* injected with varying concentrations of MBs. Scale bars, 0.5 mm. **g**, Average flow speed values of ULM-3,000v (left) and FLAME-120v (right). The solid curves are fitted through a linear regression model. Whiskers, 75% and 25%.

**Fig. 3.**
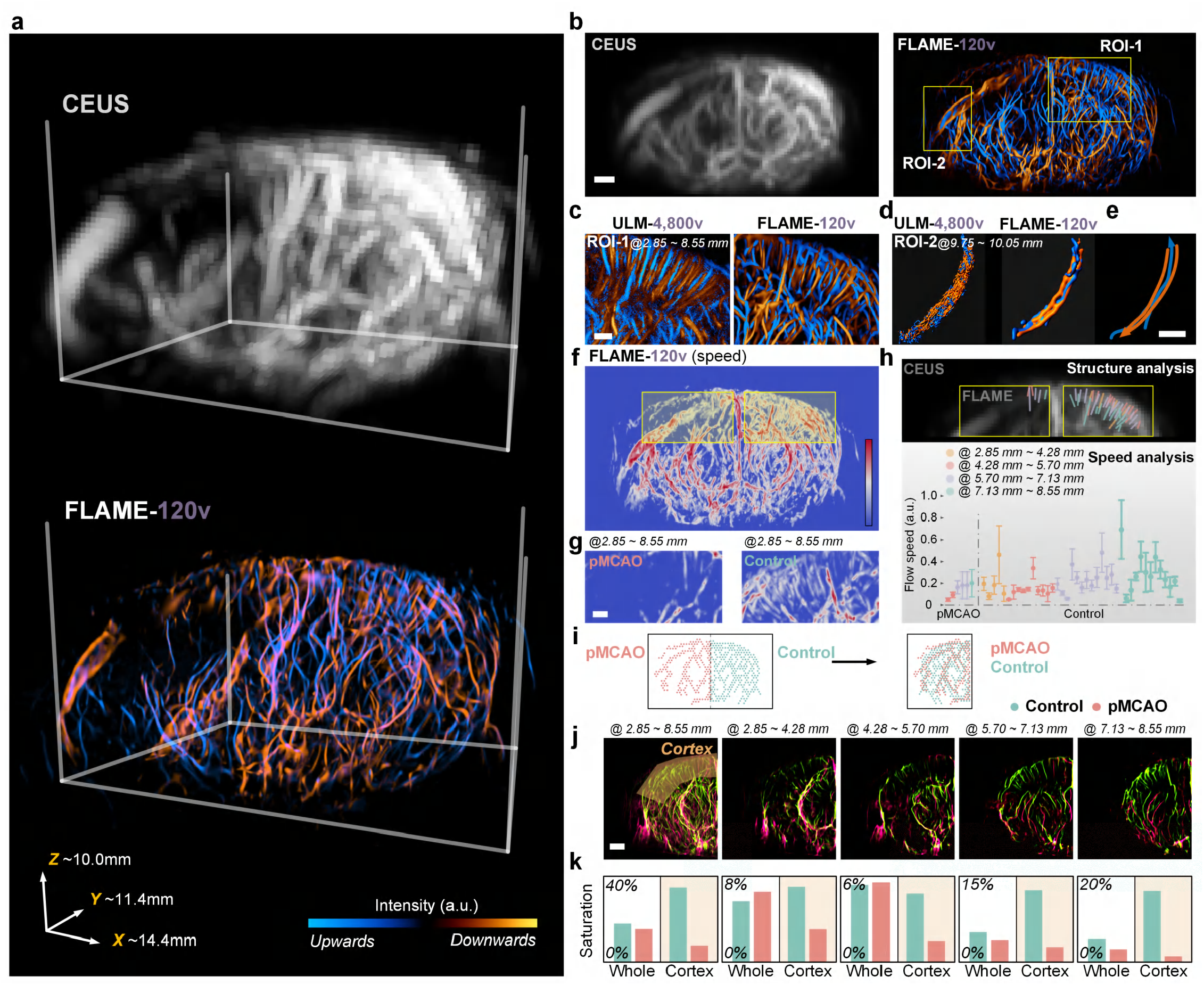
FLAME enables characterization of stroke in mouse-brain microvasculature. **a**, 3D rendering of a whole-brain microvascular network in a mouse with intact scalp and skull recorded by CEUS (top) and FLAME-120v (bottom). Volume size is labeled at the bottom left corner. **b**, Maximum intensity projection of the microvasculature in (A) under CEUS (gray, left) and FLAME-120v (right) in dual-channel. Upward and downward directions are labeled with orange and blue, respectively. Scale bars, 1 mm. **c**, Magnified views of the yellow box (ROI-1) in (**b**) reconstructed by ULM-4,800v (left) and FLAME-120v (right). Scale bars, 0.5 mm. **d**, Surface renderings of selected vessels (ROI-2) in (**b**) recorded by ULM-4,800v (left) and FLAME-120v (right). **e**, Schematic diagram for the structures and flow directions of vessels in (**d**). Scale bars, 0.5 mm. **f**, Maximum flow speed projection maps of FLAME-120v. Scale bars, 1 mm. **g**, Magnified views of the yellow box in (**f**) under the pMCAO (left) and healthy (right) hemispheres. Scale bars, 0.5 mm. **h**, Cortical vessels in CEUS cannot be extracted (top), while FLAME enables reconstructions of individual vessels for flow speed analysis (bottom). Average flow speed values of FLAME-120v under pMCAO (left) and healthy (right) hemispheres. Colors indicate different location depths (along the *y* axis) of vessels; whiskers, 75% and 25%. (**i, j**) Schematic diagram (**i**) and merged coronal views under FLAME-120v (**j**) of the pMCAO (salmon color) and healthy (turquoise color) hemispheres. Scale bars, 1 mm. **k**, Saturation rates of whole hemispheres (‘whole’) and cortical (‘cortex’) regions defined as the fraction of voxels occupied by vessels, which were calculated by the proportion of non-zero pixels in the binary masks of FLAME reconstructions against overall pixel numbers in the cortex and whole brain, respectively.

Unlike ULM, benefiting from its group-level reconstruction strategy, FLAME produced high-quality reconstructions with different experimental conditions, across different data acquisition durations (**Extended Data Fig. 3b, Extended Data Fig. 4a-4c**), as well as 10-fold variations in MB densities (**Extended Data Fig. 5, Extended Data Fig. 6c**) and frame rates (**Extended Data Fig. 6a-6b**). To assess the reconstruction convergence speed, we analyzed the pixel saturation rate and structural similarity index (SSIM)^36^ of the FLAME results against ULM-4,800v as the reference (**Fig. 2d**). We can observe that FLAME exhibited a sharp increase in SSIM within the initial 30 volumes and reached a plateau in saturation rate by 120 volumes (**Fig. 2d**). In contrast, ULM showed a much slower SSIM increase even beyond 1,800 frames. These quantitative results further demonstrate that FLAME achieves higher throughput than ULM and outperforms other fluctuation-based super-resolution methods (**Extended Data Figs. 7**-**8**). With an imaging frame rate up to 1.2 kHz (Methods, **Extended Data Fig. 2**), FLAME achieves a super-resolution volume rate of ∼40 Hz, which has not been demonstrated by 3D-ULM methods.

In flow speed measurement, the flow maps derived from FLAME-120v closely matched ULM-4,800v, with nearly identical local distributions (**Fig. 2e, Extended Data Fig. 4d**-**4e**). We further assessed the accuracy and dynamic range of the flow measurement on a custom-designed phantom at varying flow speeds (**Fig. 2f**). It is observed that FLAME (using 120 volumes) exhibited a strong linear relationship with ground-truth over a large flow range, whereas ULM (using 3,000 volumes) failed under both low and high flow regimes (**Fig. 2g, Extended Data Fig. 3c**). We would like to note that, although FLAME cannot provide absolute flow quantification, it can provide a useful flow index that reliably reflects the hemodynamic change, similar to laser speckle imaging^33^.

### Super-resolution characterization of stroke in mice

Characterization of whole-brain hemodynamics is critical for treating post-stroke ischemic injury. Here, FLAME offers non-invasive estimations of brain perfusion in the stroke mouse model induced by permanent middle cerebral artery occlusion (pMCAO), without compromising the temporal resolution or penetration depth of traditional CEUS (Methods, **Fig. 3a, 3b, Supplementary Video 3**). Although ULM ^16^ can similarly provide high-resolution microvascular maps of the stroke brain by using 4,800 volumes (**Fig. 3c**), FLAME offers high-fidelity images across a broader range of flow speeds, by using only 120 volumes (**Fig. 3c**-**3e, Supplementary Fig. 1**). For example, ROI-1 covers a FOV over ∼73 mm^3^ in the healthy hemisphere (**Fig. 3c**), where both ULM-4,800v and FLAME-120v yield comparable microvascular structures. For ROI-2 near the pMCAO region (**Fig. 3d, 3e**), FLAME-120v shows three large blood vessels with well-defined boundaries and clearly-distinguished fast directional flow, likely reflecting stroke-induced occlusion^37^. However, the ULM reconstructions appear less discernible for these large vessels, because the relatively high MB density hinders reliable localization and tracking of individual bubbles. To examine functional changes due to the ischemic injury, we compared the flow speed maps between the two hemispheres (**Fig. 3f**). Two representative ROIs in the cortex were chosen from the stroke and control regions, revealing distinct flow distributions of diving vessels in the cortex (**Fig. 3g**), while the CEUS images largely cannot resolve the individual vessels (top panel in **Fig. 3h**). In quantitative analysis, it demonstrated a significant flow-speed decrease in the diving vessel on the stroke side (**Fig. 3h**). Along the brain’s vertical axis, significant flow reductions were observed in the cortex of the stroke hemisphere within the 5.70−7.13 mm and 7.13−8.55 mm depth ranges (**Fig. 3i-3k**), which were most affected by pMCAO. By contrast, the functional difference at the hemispherical scale appeared less pronounced due to the spatial averaging (**Fig. 3k**), which highlights the importance of single-vessel resolution enhancement by FLAME. Together, these examples validate FLAME’s capacity to provide high-resolution vasculature and robust flow quantification across varying vessel sizes, depths, and flow speeds.

### Instant capturing of brain-wide acute responses

FLAME can facilitate rapid volumetric recording of brain-wide hemodynamics at microscopic scale, allowing precise evaluation of global and localized responses. In the global response study, we examined age-dependent brain response to a single dose of epinephrine by using time-lapse FLAME in young and aged mice under otherwise identical conditions (**Fig. 4a-4d, Supplementary Video 4**). To improve temporal continuity in dynamic monitoring, we applied a rolling-window operation to the FLAME results (reconstructed from every 30 CEUS volumes) by averaging 4 consecutive volumes with a one-volume step. The hemodynamic responses in blood perfusion (**Fig. 4b**) and flow speed (**Fig. 4c**) of young and aged mice exhibit different patterns. The kymographs (**Fig. 4d**) extracted along the yellow regions in **Fig. 4c** reveal the changes started at ∼30 seconds when the epinephrine was injected. Quantitative comparisons (**Fig. 4e**) indicate that the perfusion changes were primarily driven by microvessels rather than macrovessels. Young mice show a fast and transient reaction, whereas aged mice display a prolonged response without recovery in the observation window. These dynamics were further characterized by the saturation rates, defined as the ratio of signal-occupied pixels to the total number of pixels within the selected region, and flow speeds over time (**Fig. 4f-4g**). The flow speed distributions at three representative time points (**Fig. 4h**) confirmed a transient flow speed increase followed by normalization in young mice but remained high in aged mice.

**Fig. 4.**
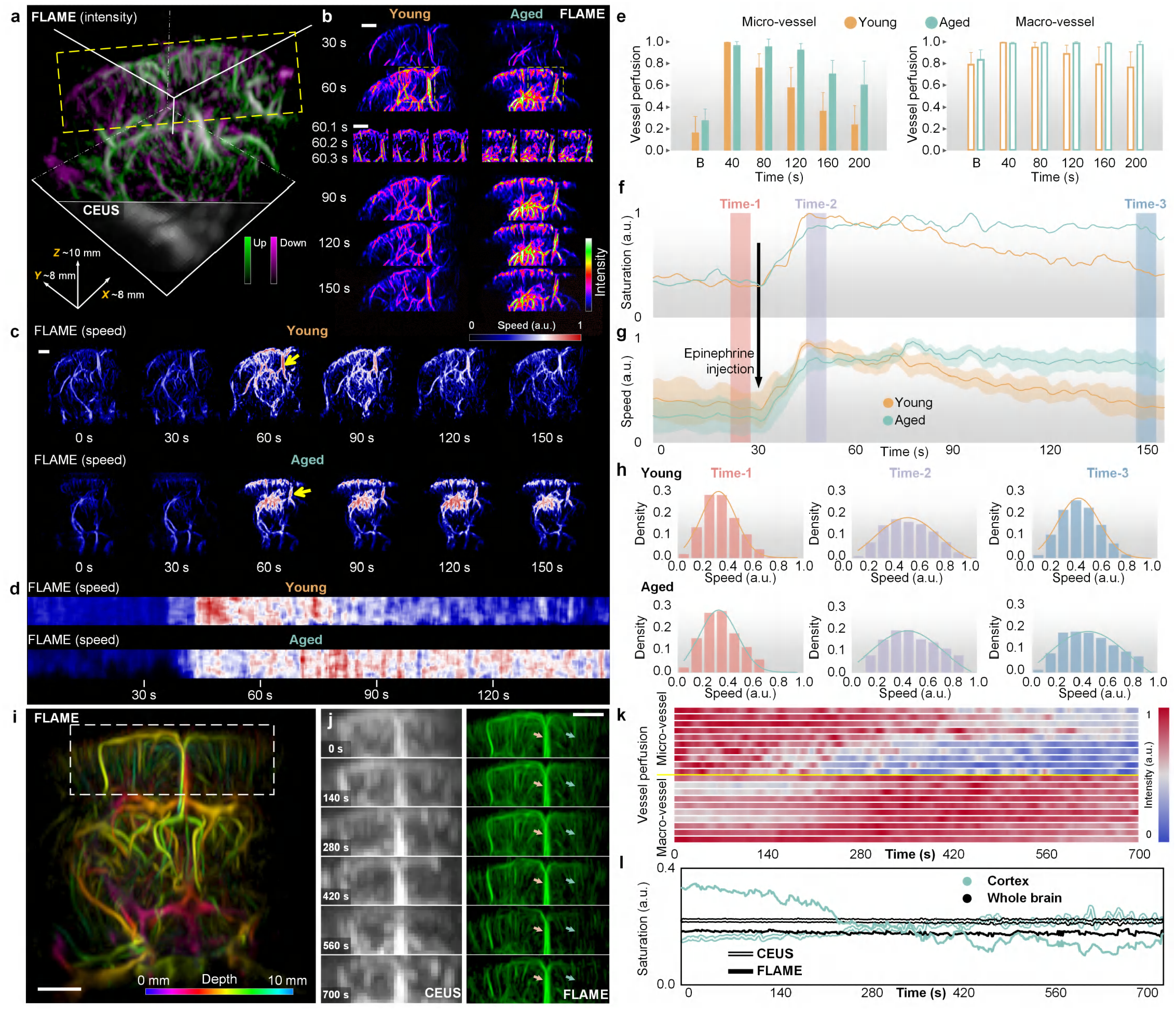
FLAME reveals the fast mouse-brain microvascular dynamics. **a**, 3D rendering of cerebral microvasculature in a living mouse with intact scalp and skull recorded by FLAME (top) and CEUS (bottom). Volume size is labeled at the bottom left corner. **b**, Representative montages of maximum intensity projections from the yellow dashed box in (**a**) at 6 time points, showcasing cerebral microvasculature in response to one dose of epinephrine (injection at 30 s) in young (left) and aged (right) mice. Insert: Magnified views of 3 consecutive time points from the yellow dashed box. Recorded by FLAME in 10 volumes/s using a 120-volume (0.4 s) window with a 30-volume (0.1 s) step. Scale bars, 1 mm. **c**, Full-view montages of maximum flow speed projections from (**a**) at five time points in young (top) and aged (bottom) mice. Scale bars, 1 mm. **d**, Kymographs derived from vessels pointed by yellow arrows in (**c**). **e**, Comparisons of the micro- and macrovessel perfusion changes in young (khaki) and aged (turquoise) mice. In this work, the perfusion is defined as the normalized average intensity values of center pixels within individual vessels. (**f, g**) Saturation rate of cortex (**f**) and average flow speed (± s.e.m.) of microvessels (**g**) in young (khaki) and aged (turquoise) mice during response to one dose of epinephrine (injection at 30 s). **h**, Histogram of speed distributions from a single microvessel in young (khaki) and aged (turquoise) mice at 3 selected time points. **i**, Color-coded 3D distributions of the microvasculature in a living hybrid transgenic mouse, Chm4 (see Methods), with a CNO (Clozapine-N-oxide) injection. Scale bars, 1 mm. **j**, Representative maximum intensity projections from the white dashed box in (**i**) under CEUS (left) and FLAME (right) at 6 time points. Scale bars, 1 mm. **k**, Comparisons of the micro-(top) and macro (bottom) -vessel perfusion changes across ∼12 minutes FLAME imaging. **l**, Saturation rate changes in the cortex (turquoise color) and whole brain (black color) under CEUS (hollow) and FLAME (solid).

Next, to assess local response in the brain (**Fig. 4i, Supplementary Video 5**), we evaluated the vascular effects of Clozapine-N-oxide (CNO) in a transgenic mouse model (Methods). Unlike epinephrine, which induces systemic responses, CNO selectively affects cortical microvessels, as confirmed by using two-photon fluorescence microscopy (**Supplementary Fig. 2**). Due to the poor resolution, CEUS fails to detect any significant microvascular changes, whereas FLAME readily resolves a localized decrease in capillary perfusion (turquoise arrows in **Fig. 4j**) while showing largely persistent perfusion in macrovessels (khaki arrows in **Fig. 4j**). Such selectivity is consistent with the fact that CNO activates engineered DREADD (Designer Receptors Exclusively Activated by Designer Drug) receptors expressed in specific vascular cell types, which regulate the local microvascular constriction without influencing larger vessels. These responses were monitored over a 12-minute observation window, revealing perfusion declines restricted to microvessels (**Fig. 4k**). Moreover, FLAME successfully delineated the cortical hemodynamic changes with high resolution and sensitivity, which were not obvious on the whole-brain level (**Fig. 4l**). Similarly, we performed hypoxia stimulation in mice, which highlighted the superior spatiotemporal resolutions of FLAME essential for monitoring acute hemodynamic changes (**Supplementary Fig. 3**).

### High-throughput *in vivo* hemodynamic imaging at whole-body scal

Super-resolution imaging of small animal models at the whole-body level is technically challenging, mostly due to the strong motion artifacts induced by the animal’s breathing. Because FLAME utilizes substantially fewer volumes than ULM, it is inherently less sensitive to motion artifacts. For whole-body mouse imaging, the animal was placed in a prone position and scanned using a 2D motor stage (**Fig. 5a, Supplementary Video 6**), acquiring 120 volumes per FOV.

**Fig. 5.**
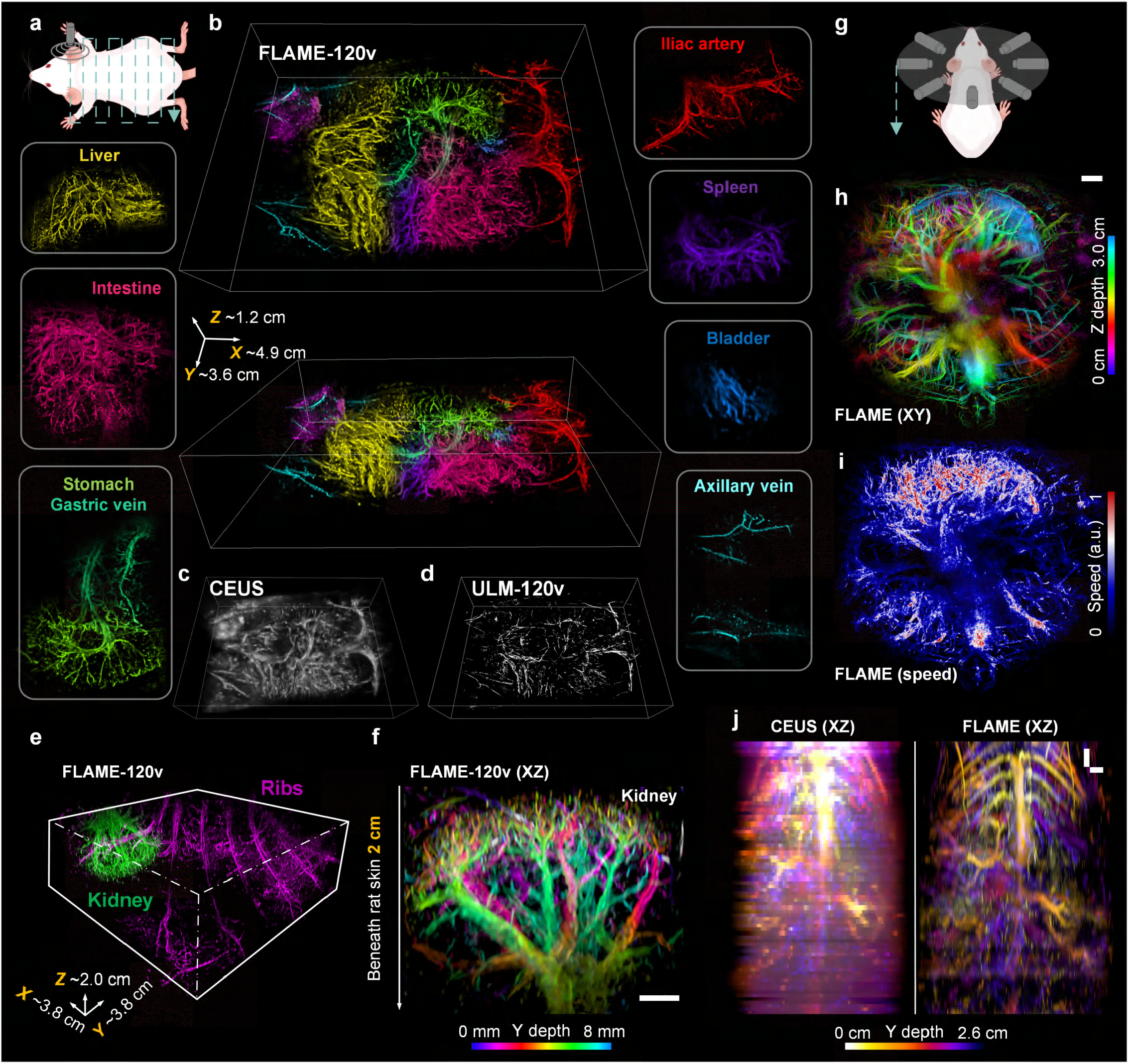
Whole-body microvascular system of rodents recorded by FLAME. **a**, Schematic of the acquisition configuration for ultrafast volumetric whole-body CEUS and FLAME imaging (details in Extended Data Fig. 2a). **b**, 3D rendering of whole-body mouse microvasculature imaged by FLAME-120v, surrounded by enlarged regions of different organs coded with different colors. Volume size is labeled at the bottom left corner. (**c, d**) 3D renderings of corresponding CEUS (**c**) and ULM-120v (**d**) results. **e**, 3D rendering of renal-region microvasculature in a living rat recorded by FLAME-120v. Volume size is labeled at the bottom left corner. **f**, Zoomed color-coded projection view from the green color (kidney) region in (**e**). Scale bar, 2 mm. **g**, Schematic of an alternative acquisition configuration for volumetric whole-body CEUS and FLAME imaging (details in Extended Data Fig. 2b). **h**, Color-coded horizontal (XY projected) view of the primary torso microvasculature in a living mouse recorded by FLAME-120v. Scale bar, 2 mm. **i**, Maximum flow speed projection map of FLAME-120v corresponding to (**h**). **j**, Color-coded vertical (XZ projected) views under CEUS (left) and FLAME-120v (right). Scale bars, 2 mm. [Schematics were created with www.figdraw.com]

The whole-body imaging was completed in under 1.5 minutes. Notably, the imaging acquisition per FOV is fast (< 30 seconds for 126 FOVs); and the more time-consuming process is the mechanical translation of the animal (> 60 seconds). As shown in **Fig. 5b**, FLAME-120v enabled comprehensive 3D visualization of the entire mouse vasculature system, delineating internal organs with high contrast and little motion artifacts. In comparison, ULM-120v (**Fig. 5c-5d**) yielded sparse and discontinuous vessel maps, particularly under high MB density and short acquisition time (**Extended Data Fig. 9a-9b**). This result is important to demonstrate that FLAME can relax the imaging constraints of tracking-based technologies, such as the MB density, data acquisition time, and imaging speed, all of which are beneficial for high-throughput applications. To evaluate FLAME’s scalability in even deeper tissues, we imaged the left kidney of a living rat (**Fig. 5e–5f, Supplementary Video 7**), revealing detailed vascular organization and branching patterns at a depth of ∼2 cm with only 120 volumes, despite the acoustic attenuation and tissue heterogeneity (**Extended Data Fig. 9c-9d**).

To test the broad applicability of FLAME on different imaging setups, we also implemented FLAME using a ring-array transducer with a 5-MHz center frequency (**Fig. 5g, Extended Data Fig. 2b, Supplementary Video 8**). FLAME on this setup yielded cross-sectional super-resolution images of dense vascular networks in the mouse torso (**Fig. 5h, 5j**), along with the corresponding flow speed maps (**Fig. 5i**). For 3D reconstruction, we translated the animal along the elevational direction, acquiring a total of 30 cross-sectional datasets, with a step size of 0.4 mm. To mitigate aliasing from the coarse elevational sampling, we applied a 4× Fourier-based upsampling (**Extended Data Fig. 10a-10c**). Quantitative resolution assessment was performed using the previously developed rolling Fourier ring correlation (rFRC)^35^ across five cross-sectional segments (**Extended Data Fig. Fig. 10d**). We observe that CEUS showed resolution degradation with depth from 346 μm to 448 μm, and FLAME maintained relatively consistent resolution from 41 μm to 57 μm throughout the imaging depth, with minimal local resolution approaching 20 μm. Together, these results demonstrate FLAME’s versatility for multi-scale high-throughput *in vivo* imaging under different imaging conditions and platforms.

## DISCUSSION

In conclusion, we demonstrate FLAME as an all-acoustic solution for fast, deep, and super-resolution hemodynamic imaging. By circumventing the need for MB localization and tracking, FLAME accelerates the imaging pipeline with much shorter acquisition time using higher MB concentration and increases the compatibility with high flow speeds and low SNR conditions. FLAME retains the large penetration depth, high temporal resolution, and high system compatibility of conventional CEUS, while achieving up to 8-fold 3D spatial resolution enhancement without the need for hardware modification. FLAME’s balanced resolution, speed, and accessibility fulfill the long-standing need for high-throughput, deep-tissue microvascular imaging with existing

US imaging setups. Executed on less than 1 gigabyte (GB) of data, FLAME completes GPU-accelerated reconstruction within half a minute using just 30 volumes. By contrast, volumetric ULM typically requires over 1 terabyte (TB) of data from 4,800 volumes and several hours to generate a single super-resolution volume. Overall, FLAME’s potential application on portable or wearable US devices could enable longitudinal *in vivo* monitoring of systemic hemodynamics in freely behaving animals and ultimately in humans. Further integration of high-throughput FLAME with therapeutic US technologies, such as US-mediated drug delivery or neuromodulation, may offer a promising strategy for real-time treatment monitoring and personalized therapy of diverse diseases^38,39^.

Further development of FLAME will progress along two major directions. Firstly, to improve the performance, several avenues are promising. (i) Multimodal integration with photoacoustic imaging will enable simultaneous super-resolution functional and molecular imaging of multiparametric tissue microenvironment beyond hemodynamics^16^. (ii) High-order cumulant analysis holds promise for estimating local MB velocities and densities. Distinct from the Bernoulli distribution assumed in fluorescence imaging^40^, the MB motion model should be carefully investigated^41^. (iii) Adaptive reconstruction using spatially variant PSFs can be applied to moderate structural distortions across the FOV. Secondly, to facilitate clinical translation, advances in real-time processing and label-free imaging are essential. Real-time visualization, combined with fully automatic and adaptive quantitative analysis, will be critical for clinical emergency care and bedside monitoring, providing timely feedback for various medical conditions. Furthermore, the relaxed requirement for MB concentration control and high-SNR conditions may expand FLAME to leverage endogenous red blood cells as intrinsic contrast sources for label-free microvascular mapping. Transitioning FLAME toward label-free US imaging will eliminate the need for exogenous contrast agents, hence streamlining clinical workflows and broadening its applicability for longitudinal and routine monitoring. Taken together, we anticipate that the continuous development of FLAME will improve the current super-resolution US imaging, offering a more flexible platform for both preclinical research and clinical practice, particularly in real-time applications where conventional high-resolution tracking-based US imaging remains challenging.

## METHODS

### FLAME framework

#### Overall workflow

FLAME reconstruction (**Fig. 1b** and **Extended Data Fig. 1**) consists of three major steps: (Step 1) Multi-domain pre-filtering for high-quality microbubble (MB) motion signal extraction; (Step 2) High-order cumulant analysis of MB-induced temporal fluctuations for resolution enhancement; (Step 3) Final post-deconvolution for maximizing spatial resolution and image quality. In Step 1.1, the collected raw IQ (in-phase and quadrature-phase) volumes contain strong tissue scattering signals on a large stationary scale. To isolate MB signals, a tissue signal filter is applied in the temporal domain using singular value decomposition (SVD). In Step 1.2, an MB direction filter is applied in the Fourier domain to separate upward and downward flow components relative to the transducer, providing directional flow information. Despite SVD filtering, residual tissue signals may persist due to large-scale in vivo motion. To address this, Step 1.3 applies a background filter in the wavelet domain to suppress leaked low-frequency components. In Step 1.4, a 3D Richardson–Lucy (RL) deconvolution is performed as a pre-deconvolution to reduce noise, eliminate clutter, and narrow the point spread function (PSF). As a key component, this step sharpens the image and enhances temporally correlated MB fluctuations without introducing noticeable artifacts, thereby improving the efficiency of subsequent high-order cumulant analysis. Step 1.5 introduces an MB density filter to further separate MB events temporally, resulting in a longer sequence with sparser point sources. Proceeding to Step 2, high-order cumulant calculations are performed along the temporal axis of each voxel, followed by brightness correction to preserve structural integrity. In Step 3.1, a 3D sparse deconvolution is applied to further refine spatial resolution and optimize volume quality. Finally, data fusion is performed to generate the final super-resolved intensity and flow speed maps. Notably, prior to the deconvolution steps, a 3D Fourier interpolation is performed to refine voxel dimensions and support higher-resolution reconstructions.

#### Stationary tissue signal filter (temporal domain)

An SVD-based clutter filter^28^ was applied to extract the MB signal from the surrounding stationary tissue signal. The full dataset was divided into local batches (typically 60 volumes per batch in this work), and SVD filtering was applied individually to each batch. In each batch, the collected ultrasound (US) signal is written as follows:

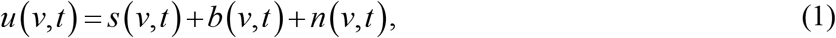

where *v* represents the volumetric spatial coordinate and *t* denotes the sequential recording along time. *s* is the stationary tissue signal, which is highly coherent both in space and time, occupying the low-rank subspaces with high singular values; *b* is the MB flow with rapid fluctuations, residing in the high-rank subspaces with low singular values; and *n* is the noise components. After SVD processing, the *u* in Eq. (1) can be rewritten as:

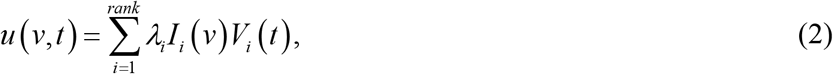

where *rank* represents the number of singular vectors; and *I* and *V* denote decomposed spatial images and time signals. Based on the assumption that tissue signal is gathered in the first singular vectors, the SVD filter is performed to remove the low-rank singular vectors from the raw signal.

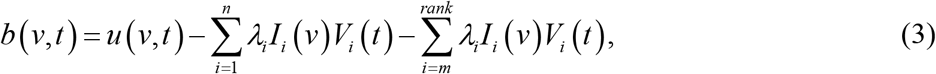

where *m* and *n* represent the estimated singular value cutoff thresholds (equivalently, as a bandpass filter). Particularly, in the whole-body imaging application, we adopt a block-wise adaptive SVD filter^29^ to optimize the separation of tissue and MB signals with varying flow speeds across different local regions.

#### MB direction filter (Fourier domain)

To extract the flow direction information, we applied a simplified MB direction filter^30^ to the SVD-filtered data. Because the Doppler frequency shift is assumed to be proportional to the MB’s velocity along the US beam direction, with positive frequencies indicating movement toward the transducer and negative frequencies indicating movement away, the MB signal within a certain direction can be extracted by a bidirectional filter (setting the first and second halves to zeros, respectively) in the Fourier domain. We applied a 1D Fourier transform along the temporal axis (*xyz* reshaped as one spatial axis) and reassembled the filtered data with positive and negative directions into dual-direction volumes. Before the Fourier transform, a Hamming window was applied to avoid artifacts induced by the fast Fourier transform. After the directional filter, the complex value data is converted to their magnitudes, and the subsequent steps are executed individually on these two volumes.

#### Background filter (wavelet domain)

We also included a background subtraction step in the wavelet domain to remove the low-frequency baseline signal and residual tissue background. The background is iteratively estimated from the lowest-frequency wavelet bands using 2D Daubechies-6 filters up to the 7th level^10^. To preserve dim MB signals, we apply an inverse wavelet transform, compare the result with half of the square root of the input image, and merge them by keeping the minimum pixel values. This off-peak background estimation is typically repeated three times, and the estimated result is subtracted from the input image.

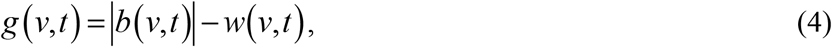

where |.| is the modulus calculation, and *w* represents the estimated background.

#### Pre-deconvolution (spatial domain)

A pre-deconvolution is used to simultaneously reduce the noise, reject the out-of-focus signals, and improve the resolution. Before this step, we execute a 3D Fourier interpolation to fulfill the subsequent spatial resolution improvements, and after that, the voxel size is 2-fold finer (for example, from 120 *μ*m to 60 *μ*m). To do so, the resolution enhancement ability of the temporal correlation cumulant can be used more efficiently. We employ an accelerated 3D Richardson-Lucy (RL) deconvolution without any regularizations in this work^11^:

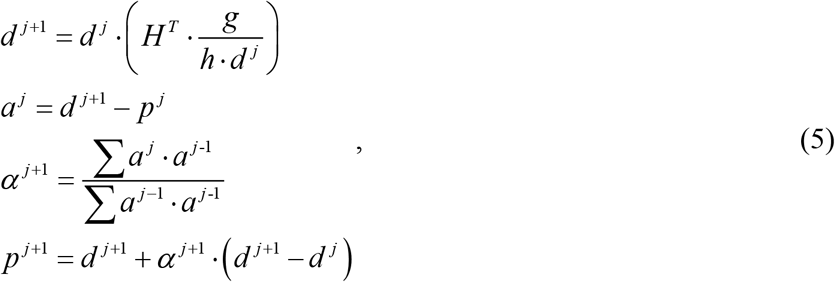

where *p* ^*n*+1^ is the image after *n*+1 iterations; *g* is the input image; and *H* is the PSF. The adaptive acceleration factor *α* represents the length of an iteration step, which can be estimated directly from the experimental results^42^. We used a 3D Gaussian kernel with of theoretical 3D resolution to approximate the US PSF. Without needing localization, our pre-deconvolution relaxes the requirement for precise measurement of the system transfer function.

#### MB density filter (temporal domain)

We employ the Haar wavelet kernel (HAWK) analysis^31^ to decompose the deconvoluted high-quality data into a longer sequence with reduced MB density per volume. Each pixel is independently treated as a time sequence and convoluted with multi-level Harr wavelet kernels. Then the resulting sequences are concatenated to produce the final extended sequence.

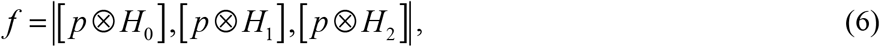

where *H* represents the Haar kernels, and *p* is the data after pre-deconvolution. MBs moving out of the local field of view (FOV) may produce negative values, which are converted to absolute values. Typically, we apply three levels of Haar kernels to generate a 3-fold longer sequence, i.e., from 30 volumes to virtual 90 volumes.

#### High-order fluctuation analysis

After executing multi-domain pre-filters, the sample can be simply regarded as composed of *N* single independently moving MBs located at position *v*_*k*_.

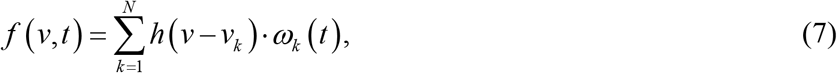

where *h* and *ω* represent the narrowed PSF after pre-deconvolution, and the time-dependent function of MB motion. Calculation of the second-order auto-correlation, *G* ^2^, with zero time-lag along time is written as follows:

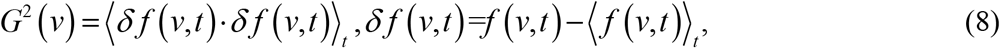

where ⟨ · ⟨ _*t*_ denotes the time averaging. Then, if expanding the expression of *G* ^2^, we can obtain the following formula.

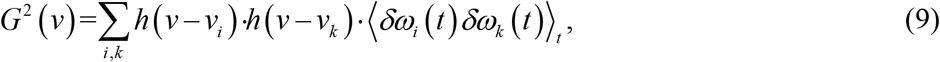

The flow of MBs along the vessels results in correlated fluctuations in neighboring pixels located along the streamlines and uncorrelated fluctuations in different vessels. As a result, assuming that each MB is moving individually, the cross-correlation terms in this equation can be regarded as zero when *i* ≠ *k*. Therefore, such second-order temporal cumulant can be written as the simple sum of the squared PSF weighted by the corresponding constant brightness *γ*.

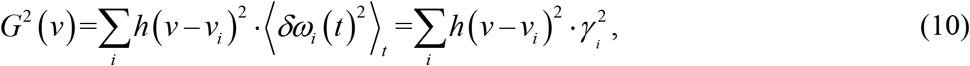

where the *γ* is the average MB scattering *d* values. Here, the width of the resulting PSF is reduced by a factor of 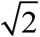. Moreover, by isolating sparse MB emitters through multi-domain pre-filters, high-contrast fluctuations with appropriate blinking densities enable the application of high-order cumulant analysis to suppress low-order correlations and maximize spatial resolutions:

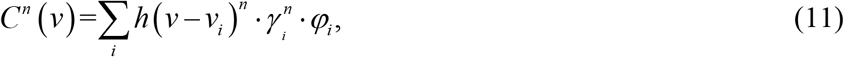

where *φ* represents the high-order correlation-weighted value. The 6^th^-order cumulant *C* ^*6*^ can be derived from the auto-correlation functions *G*, using the recursive formula from ref. ^43^.

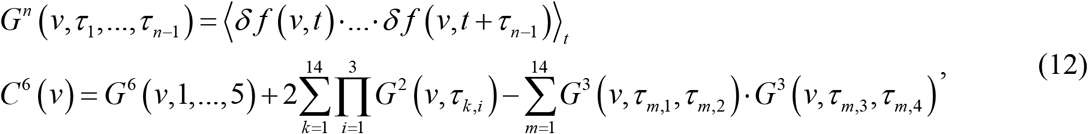

where *τ* represents the time lag. We typically used 30 volumes for the 6th-order cumulant calculation, attaining 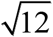-fold resolution improvements (combined with the pre-deconvolution) across all three dimensions.

#### Brightness correction

When applying the high-order statistics, we found that the dynamic range of the reconstructed image increases, and the imaged structures appear grainy. Different from fluorescence microscopy (fluorophores are static and truly blinking), CEUS imaging is a dynamic system (MBs flow along the vessels), and thus the expanded dynamic range of FLAME is moderated compared to fluorescence imaging. To fully eliminate the intensity nonlinearity caused by the motion correlation heterogeneity (*φ*), we adopt a linear dynamic range compression^32^ method with the corresponding second-order results as a target for brightness correction without canceling the resolution improvements.

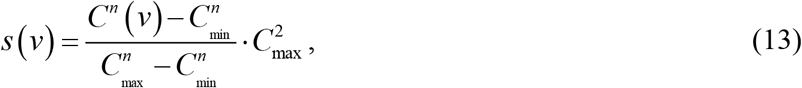

where *C* ^*n*^ and *C* ^2^ denote the pixel intensity in n^th^ order and second order cumulant results, respectively. This intensity normalization according to the reference *C* ^2^ is performed locally in a small window that is scanned across the image in a plane-by-plane manner. The footnotes max and min represent maximum and minimum pixel intensities in the window.

#### Post-deconvolution

Finally, we extend our previously developed Sparse deconvolution^10^ as a 3D post-processing approach for optimal spatial resolution enhancement. Instead of processing each plane individually, the approach operates volumetrically in this case. Specifically, the multi-constraint deconvolution is given by:

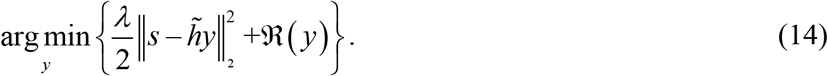

The first term on the left side is the fidelity term, representing the distance between the recovered image *x* and the result obtained with the input image *s*. 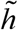 is the 6-fold improved PSF. The second term represents the continuity and sparsity constraints, which are given by:

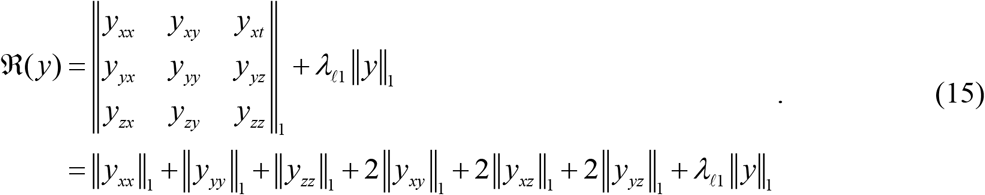

The first term is the continuity prior, and the second term is the sparsity constraint. *λ* and 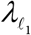 denote weight factors to balance image fidelity with sparsity. The subscript *xx* is the second-order derivation operator in the *x*-direction. To ensure sufficient Nyquist sampling criteria for ensuring maximal spatial resolution, the PSF must occupy more than 3 × 3 × 3 pixels in space, which constitutes the basis for the volumetric continuity. Moreover, the US signals are blurred by diffraction and accompanied by noise during acquisition, making the true signal inherently sparser than the recorded one. Therefore, FLAME incorporates continuity and sparsity priors to enhance resolutions and reduce artifacts. As described previously, to solve such convex optimization problem, we split it into constraint reconstruction and deconvolution subproblems (details in ref. ^10^). A 3D Fourier interpolation before this step is executed to fulfill the subsequent spatial resolution improvements, and after that, the voxel size is 4-fold finer (for example: from 60 *μ*m to 15 *μ*m).

#### Data fusion

Notably, the post-deconvolution step also provides the basis for correcting the 6^th^ power brightness caused by the 6^th^ order cumulant, without cancelling the improvement in resolution. Accordingly, we applied a 1/6 root at first to directly linearize the brightness, followed by a convolution of 8-fold narrowed 3D Gaussian PSF, comprising a 6-fold increase from the 6th-order cumulant combined with post-deconvolution plus an additional ∼1.4-fold from pre-deconvolution^11^.

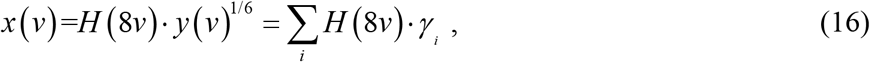

Then, we perform average and normalized standard deviation followed by percentile normalization for generating high-quality FLAME intensity and flow speed reconstructions, respectively.

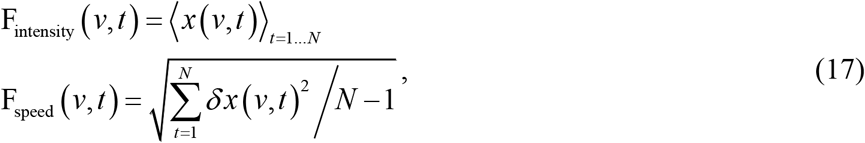

We typically set *N* = 4 when using 120 volumes to create high-quality one FLAME intensity and corresponding flow speed reconstructions. Although 30 volumes are sufficient for super-resolution reconstruction, this short temporal window for MB perfusion accumulation results in sparse signals. To preserve structural integrity and temporal resolution, dynamic intensity and speed FLAME reconstructions are generated using a rolling window of 120 volumes with a 30-volume step. Inspired by temporal laser speckle imaging^33^, using laser-induced speckle changes to measure the flow speed, we realized that MBs follow physiologically relevant blood flow trajectories, constantly altering their relative positions and directions, which is similar to the statistical blinking of laser speckles. Groups of MBs flowing through large vessels exhibit high speeds and complex dynamics, resulting in low correlation, while those in capillary beds move more slowly and steadily, maintaining high correlation. These variations create detectable changes in spatiotemporal flow dynamics, especially after super-resolution reconstruction with a short temporal window. To capture these dynamics, we use a normalized standard deviation (a.k.a., speckle temporal contrast) to map temporal MB changes, reflecting relative flow speed in local regions. Notably, further measurements and calibrations are required to establish a precise relationship between the MB velocity and our FLAME speed map.

#### Corrections of system drift and animal local motion

For large-scale motion data, we obtain the displacement matrices by executing intensity-based 3D rigid registration (translation transformation) on the averaged (per 30 volumes) raw IQ data with the previous volume as a reference, and these displacement matrices are then interpolated and applied to the results before Step 3. For small-scale motion data, we directly apply intensity-based 3D elastic registration (affine transformation) before Step 3 to compensate for the local motions.

### Volumetric US imaging configurations

#### General description

The imaging systems and system characterizations are shown in **Extended Data Fig. 2**. A low-frequency 2D hemispherical transducer array (**Extended Data Fig. 2a**) (Imasonics, France) operating at 4 MHz is used for volumetric imaging^16^, providing a balanced penetration depth through the intact skull and tissue as well as spatial resolutions within the FOV, when compared to conventional high-frequency matrix-array-based systems. The T/R bandwidth (-6dB) is 45%, and the curvature is 40 mm. The 2D hemispherical array achieves a diffraction-limited spatial resolution of 440 μm axially and 270 μm laterally for US imaging. To expand the FOV for covering full-brain or whole-body imaging, a 2D motorized translation stage facilitates raster-scanning of the animal holder. For brain imaging, all animal experiments were conducted with mice in a supine position. During whole-body imaging, mice were imaged in the prone position.

The 2D cross-sectional imaging (**Extended Data Fig. 2b**) was acquired by a ring array-based system^44^. The customized full-ring transducer array (Imasonics, France) combines two half-rings, each with 256 elements (512 in total), 4-cm radius, 5 MHz center frequency, and two-way bandwidth (-6dB) of 60%. The elevational curvature of each element is 37mm. The ring array system achieves 200-μm radial resolution and 1-mm elevational resolution. During imaging, the anesthetized animal was fixed onto a custom-built, acoustically transparent animal holder to minimize motion. The portion of the body below the neck was immersed in a temperature-regulated warm water bath. The animal holder was then placed at the center of the ring array and raster scanned using a motorized linear actuator. A total of 30 positions (step size = 0.4mm) were scanned in an elevational direction.

#### Ultrafast US imaging scheme

For the 2D hemispherical array, to balance the imaging quality and frame rate, the IQ US data were acquired using a synthetic aperture method. The volumetric imaging was performed with a 15-element synthetic transmission aperture. Each sub-aperture IQ dataset was beamformed into a volume using a 3D delay-and-sum (DAS) algorithm. For the ring array, we employed a round-trip imaging scheme utilizing 20-angle transmissions for synthetic aperture imaging. The acquired data was reconstructed using a 2D DAS beamforming algorithm. Employing a 40-mm radius hemispherical array and 2 cm imaging depth usually requires at least a waiting time of approximately 2×60 mm/(1500 m/s) ≈ 80 μs for acoustic transmission and receiving. Unlike traditional round-trip transmission schemes, we implement a multiplexed transmission strategy by sequentially emitting three transmission waves with a fixed interval of 20 μs. This interval is selected based on the maximum time-of-flight within the field of view. Although acoustic echoes from each transmission can overlap with subsequent waves, the resulting interference exhibits a relatively stable spatiotemporal pattern. Such interference can be effectively suppressed using temporal and SVD filtering, analogous to the removal of stationary tissue signals in dynamic imaging. This approach significantly increases the effective imaging frame rate (>1.2kHz) and enables faster volumetric imaging without sacrificing image fidelity.

To accelerate data throughput during ultrasound acquisition, we developed a “ping-pong” saving scheme that enables asynchronous data writing (**Extended Data Fig. 2c**). In conventional synchronous saving, the system must wait for both data acquisition, transfer from the Verasonics system, and data writing to complete before initiating the next acquisition. This introduces significant idle time. In contrast, the “ping-pong” saving strategy allows data acquisition to resume immediately once the data buffer transfer is complete. Meanwhile, data saving proceeds asynchronously in parallel. To avoid buffer overwriting, we cyclically allocate multiple memory buffers (*N* buffers in total), alternating between acquisition and saving tasks. This overlapping of the acquisition and saving process reduces idle time and enhances efficiency. Assuming *N* reusable buffers as well as equivalent data acquisition and saving time, the theoretical saving improvement is approximately (*N*-1)/2*N*. For instance, with 4 buffers, the time reduction approaches 37.5%, and with larger *N*, the gain converges toward 50%. This approach effectively increases the acquisition-saving throughput without altering acquisition time or Verasonics control logic.

To validate the effectiveness of our high-speed imaging scheme, we performed a phantom experiment using a silicone tube perfused with diluted MBs to mimic blood flow in a vessel (**Extended Data Fig. 2d**). We continuously acquired data over a 50-second period while deliberately altering the flow speed to simulate acute hemodynamic changes. Specifically, the flow was set to a “low speed” for the first 10 seconds, increased to a “high speed” over 20 seconds, and then reverted to “low speed” for the final 10 seconds. This setup allowed us to assess the system’s sensitivity to dynamic flow transitions. The resulting flow measurements clearly captured the abrupt changes, confirming the scheme’s ability to resolve transient flow dynamics. For each saved data batch, a precise time stamp was recorded, facilitating accurate temporal tracking. In total, over 60,000 frames were acquired, corresponding to an effective frame rate of up to 1.2 kHz.

### Animal preparations

#### Animals protocol

Transgenic adult B6 mice (2-3 months) were used for Chrm4 experiments. Adult SD rats (2-3 months) were used for kidney imaging experiments. Adult CD-1 mice (2-3 months old for regular group, 18 months old for aging group) were used for all rest experiments. The sex of the animals was randomly chosen.

#### Surgical procedure and preparation for imaging

Mice/rats were initially anesthetized with 5% isoflurane in 100% air within an induction chamber until the loss of the righting reflex was observed. The mouse/rat was positioned on a stereotaxic frame, with the head secured using ear bars to induce flexion. Mice/rats were first anesthetized with 1.5% isoflurane and placed on a heating pad. The scalp or body region of interest was shaved, and residual hair was removed using a depilatory cream. The skin was then thoroughly disinfected with alternating applications of 70% ethanol.

#### MB contrast agent injections

For contrast-enhanced imaging, diluted gas-filled MBs with a diameter of 0.99+/-0.56 μm (stock concentration: 1.00+/-0.20×10^8^ bubbles/mL, Revvity, USA) were intravenously injected via the tail vein with a micro pump using a 30-gauge insulin syringe. At the beginning of imaging, 10 μL MBs were initially injected into the tail vein. During the imaging session, anesthesia was maintained using 1.5% isoflurane via inhalation and 10 μL MBs were injected every minute. The total volume of MBs injected was less than 150 µL for mice and 1 mL for rats. To prevent MB separation, gentle agitation was maintained using a magnetic stirring setup, with one magnet placed inside the syringe and another externally. If the MB suspension became visually clear, indicating depletion, the syringe was replaced at least once during the session.

#### High versus low MB concentration for *in vivo* imaging

To investigate the effects of MB concentration on vascular imaging, we conducted *in vivo* studies using two MB dilution conditions in the same mouse. High-concentration MBs were prepared by diluting the stock solution 5-fold, while low-concentration MBs were diluted 30-fold, with equal injection volumes for both conditions. These conditions were designed to evaluate the trade-off between MB density and imaging fidelity. For ULM, high MB concentrations can cause spatial overlap of bubbles, leading to localization errors, false vessel structures, and resolution degradation.

#### Stroke animal preparation

A permanent middle cerebral artery occlusion (pMCAO) model was established by modifying our previously reported transient MCAO procedure^45^. Mice were anesthetized and intubated using a 20-gauge IC catheter, then maintained under 1.5% isoflurane in a gas mixture of 30% oxygen and 70% nitrogen. Rectal temperature was controlled at 37.0 ± 0.5 °C using a heat lamp. A midline neck incision was made to expose and ligate the right common carotid artery (CCA) with 4.0 silk suture. To access the MCA, a small incision was made between the right eye and ear, and the temporalis muscle was retracted. A cranial window was created by thinning the skull with an electric drill until the inner bone layer was fractured, and the bone fragments were gently removed to expose the MCA. Permanent occlusion was achieved via coagulation, with visual confirmation of perfusion loss. All incisions were closed with interrupted nylon sutures.

#### Epinephrine injection

An epinephrine challenge model was established for investigating the acute whole-brain hemodynamics change. The femoral artery and vein were catheterized for continuous monitoring and drug delivery. Systemic arterial blood pressure and core temperature were continuously recorded using PowerLab 8/35 (AD Instruments, New Zealand). Following baseline acquisition of physiological parameters and arterial blood gases, epinephrine was administered intravenously through the femoral vein, as a single bolus (100 μL of 40 μg/mL). Resuscitation doses were selected based on current American Heart Association (Dallas, Texas) guidelines for cardiac arrest management and adjusted for mice using a logarithmic scaling method that accounts for species-specific metabolic rate and body weight^46^.

#### Transgenic mouse

A transgenic model was established for investigating the chemical modulation effect on whole-brain hemodynamics. To generate pericyte-specific Designer Receptors Exclusively Activated by Designer Drugs (DREADD) mice^47^, hM4Di-DREADD mice were crossbred with Atp13a5-CreER mice, yielding Atp13a5-CreER-CAG-hM4Di offspring. To induce Cre recombination, mice received daily intraperitoneal injections of 1% tamoxifen for 5 consecutive days. Following a recovery period, cranial windows were surgically implanted, and mice were allowed to recover for 3 days before imaging. During imaging sessions, Clozapine-N-oxide (CNO, 1 mg/kg) was administered intraperitoneally after baseline acquisition to inhibit pericyte activity. CNO-induced hemodynamic changes were predominantly observed in microvessels, with minimal effects on major cerebral vessels, consistent with the pericyte-specific targeting of the transgene verified by two-photon imaging (**Supplementary Fig. 2**).

#### Hypoxia treatment

A hypoxia challenge was conducted in anesthetized mice maintained under 1% isoflurane at a flow rate of 1.5 L/min. Mice were exposed to alternating normoxic (21% O_2_ / 79% N_2_) and hypoxic (2% O_2_ / 98% N_2_) conditions. Each cycle consisted of a 1-minute normoxia baseline, 30 seconds of hypoxia, and a 6-minute recovery period under normoxia.

### Two-photon imaging

*In vivo* two-photon imaging was performed with a customized two-photon fluorescence microscope (Leica SP8 2Photon DIVE), controlling intensity with an acoustic-optical modulator (AOM). A stable, transparent cranial window was made for imaging of the brain. Carefully removing a portion of the skull and replacing it with a cover membrane, allowing repeated optical imaging. Mice were habituated for experimental handling for at least one week before imaging. Mice were head-fixed and anesthetized (by 1% isoflurane in O_2_) during imaging. Dextran-FITC dye was used (I.V. injection), and ∼820 nm excitation wavelength was used for excitation. CNO-induced vascular activity (I.P. injection) was recorded continuously at 30 seconds per frame (**Supplementary Fig. 2**).

### ULM reconstruction and visualization

To model the PSF of the system, we manually identified isolated MB signals and fitted them to a multivariate Gaussian function as an empirical PSF. The filtered MB images were interpolated to a pixel size one-fifth of the original. Each interpolated frame underwent 3D/2D normalized cross-correlation with the empirically determined PSF of the imaging system and localized the center positions of MBs^16^. A threshold is applied to the intensity map and the cross-correlation map to remove pixels with relative intensity below 0.1 and correlation coefficient values below 0.45, thereby suppressing noise and retaining high-confidence single MB images. Using the u-track3D^48^ algorithm for MB tracking, the linear assignment algorithm was implemented to link localized MB positions across frames. A maximum pairing/linking search radius of 5 pixels was assigned, and a linear motion Kalman filter model was applied to restrict the maximum linking angle to 45 degrees, ensuring robust trajectory estimation. Based on the reconstructed MB trajectories, flow direction, and velocity maps were computed, providing functional imaging information on blood flow dynamics.

### Quantitative analysis

#### Simulation

The 3D simulation modeled multiple intertwined blood vessels in close proximity that cannot be resolved individually in the envelope image. Each volume produced flickering ground-truth images through a time-continuous Markovian process^49^, simulating MB random motion. From this process, we generated a continuous volume image sequence, which was then convolved with a 3D Gaussian PSF followed by Gaussian readout noise injections.

#### SSIM

The structural similarity (SSIM)^36^ is defined as

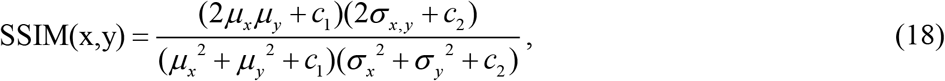

where *x* represents the reference, and *y* denotes the corresponding SR constructions. *μ* _*x*_ and *μ* _*y*_ are the averages of *x* and *y*; *σ* _*x, y*_ is the covariance of *x* and *y*; *σ* ^2^ are the variances; and *c*_1_, *c*_2_ are the variables used to stabilize the division with a small denominator.

#### FSC and rFRC

The calculations of FSC^34^ resolution and rFRC^35^ resolution map require two independent frames of identical content under the same imaging conditions. These two frames were generated by splitting the raw image sequence into two image subsets and reconstructing them independently.

#### Saturation analysis

The saturation rate was defined as the ratio of signal-occupied pixels and the total number of pixels (or long-accumulated signal-occupied pixels in **Fig. 2d**) within a selected region^50^. Maximum intensity projection (MIP) images along the y-axis were generated from the reconstructed volumes and segmented using hard thresholding. Five regions of interest (ROIs) were randomly selected within the interrogation region, and saturation was quantified as the percentage of high-intensity pixels in each ROI. The final saturation value was obtained by averaging across all ROIs.

#### Flow speed benchmarking

The central lumen regions of the vessels in the MIP view of a custom-designed *in vitro* tube phantom were manually delineated, and flow speeds were quantified under varying MB injection rates.

The experiment was repeated 25 times. Linear regression analyses were then performed on the average flow velocities within their respective applicable ranges (ULM: 0.07–0.15 μL/min; FLAME: 0.01–0.40 μL/min).

#### Flow speed analysis

For the pMCAO experiments, the imaging volume was divided into 8 slices along the y-axis. Flow speed analysis was performed on MIP views of the 4 centrally located slices with the highest vascular density. All visible vessels within these slices were delineated, and flow speeds were quantified for each vascular segment. For the epinephrine injection experiments, MIP was applied to the entire volume, and 10 representative vascular segments were randomly selected for flow speed analysis.

#### Vessel perfusion analysis

Perfusion was defined as the average intensity of the central pixels within each individual vessel, normalized to a range of 0 to 1. Microvessels (<100 μm in diameter) and macrovessels (>100 μm in diameter) were segmented from MIP views, and perfusion values were quantified for each segmented vessel. For the epinephrine injection experiments, five vessels of each type were randomly selected for analysis. For the CNO treatment experiments, ten vessels were analyzed.

#### Vessel diameter analysis

Representative vascular segments were selected from MIP view and manually delineated. Vessel diameters were quantified using a previously described method^51^, in which the full width at half maximum (FWHM) of the intensity profile was measured for each segment. Five random locations along each vessel were defined, and diameter changes at these positions were tracked over time.

### Image rendering and processing

The custom-made ‘orange-blue’ colormap was applied to render upward and downward blood flow directions in Fig. 2b, Fig. 2c, Fig. 3b, Fig. 3c, and Supplementary Fig. 3a. The custom-made ‘ipic-Phase’ colormap was used to color code the flow speed in Fig. 4c, Fig. 4d, and Fig. 5d. These colormaps are available at https://github.com/SR-Wiki/FLAMEm. The custom-made ‘sJet’ colormap was used to color-code the intensity in Fig. 4b and the resolution value in Extended Data Fig. 10d, available at https://github.com/SR-Wiki/PANELm. We used the ‘biop-12colors’ colormap to color code the 3D volumes in Fig. 2a, Fig. 4i, and Fig. 5d. We used the ‘Fire’ colormap to color code the 3D volumes in Fig. 5e. The colormap ‘biop-BrightPink’ was used to color code the intensity in Fig. 2c, Fig. 3j. The colormap ‘biop-SpringGreen’ was used to color code the intensity in Fig. 2c and Fig. 3j. The colormap ‘biop-Electrcindigo’ was used to color code the intensity in Fig. 2c. The color map ‘Green Fire Blue’ was used to color-code the intensity in Supplementary Fig. 3a bottom. The colormap ‘SQUIRREL-FRC’ was used to color-code the flow speed in Fig. 2e, Fig. 2f, Fig. 3f, Fig. 3g, and Fig. 3k. All figures were prepared with MATLAB, ImageJ, Microsoft Visio, Figdraw, and OriginPro; and videos were all produced with Imaris, Blender, and our light-weight MATLAB framework, which is available at https://github.com/SR-Wiki/img2vid.

## Data availability

All the data that support the findings of this study are available from the corresponding author upon request.

## Code availability

The tutorials and the updating version of our FLAME reconstruction can be found at https://github.com/SR-Wiki/FLAMEm.

## Acknowledgments

We thank Yirui Xu, Rui Yao, and Jinhuan Luo for the technical support. We thank Drs. Wei Yang and Liping Feng for providing the animal models. We thank Dr. Pengfei Song for guidance on ULM. This work was supported by the National Natural Science Foundation of China (grant no. 32422052 to W. Z., grant no. 62305083 to W. Z., grant no. 32227802 to L. C. grant no. 81925022 to L.C., grant no. T2288102 to L.C.), the National Key Research and Development Program of China (grant no. 2024YFC3406600 to X. D., grant no. 2022YFC3400600 to L. C.), the Scientific Research Innovation Capability Support Project for Young Faculty (grant no. ZYGXQNJSKYCXNLZCXM-H7 to H. L.), and the Beijing Natural Science Foundation (grant no. Z20J00059 to L.C.).

## Author contributions

W.Z., H.L., and Z.H. conceived and initiated the research; W.Z. designed the reconstruction algorithm; Z.H. implemented the software; N.W. implemented the data processing pipeline of CEUS and ULM for comparison; N.W. and T.Z. developed the US imaging system; N.W. designed the experiments and collected the data; Z.H., W.Z. and N.W. analyzed the data; Z.H., X.X., and X.M. prepared the videos; J.G., Y.L., Q.L., L.Q., W.L., X.D., X.L., D.T., J.T. and L.C. participated in discussions during the development of the manuscript; W.Z., N.W., and H.L. wrote the manuscript with input from all authors; H.L., W.Z., and N.W. supervised the project. All authors participated in the discussions and data interpretation.

## EXTENDED DATA FIGURES

**Extended Data Fig. 1.**
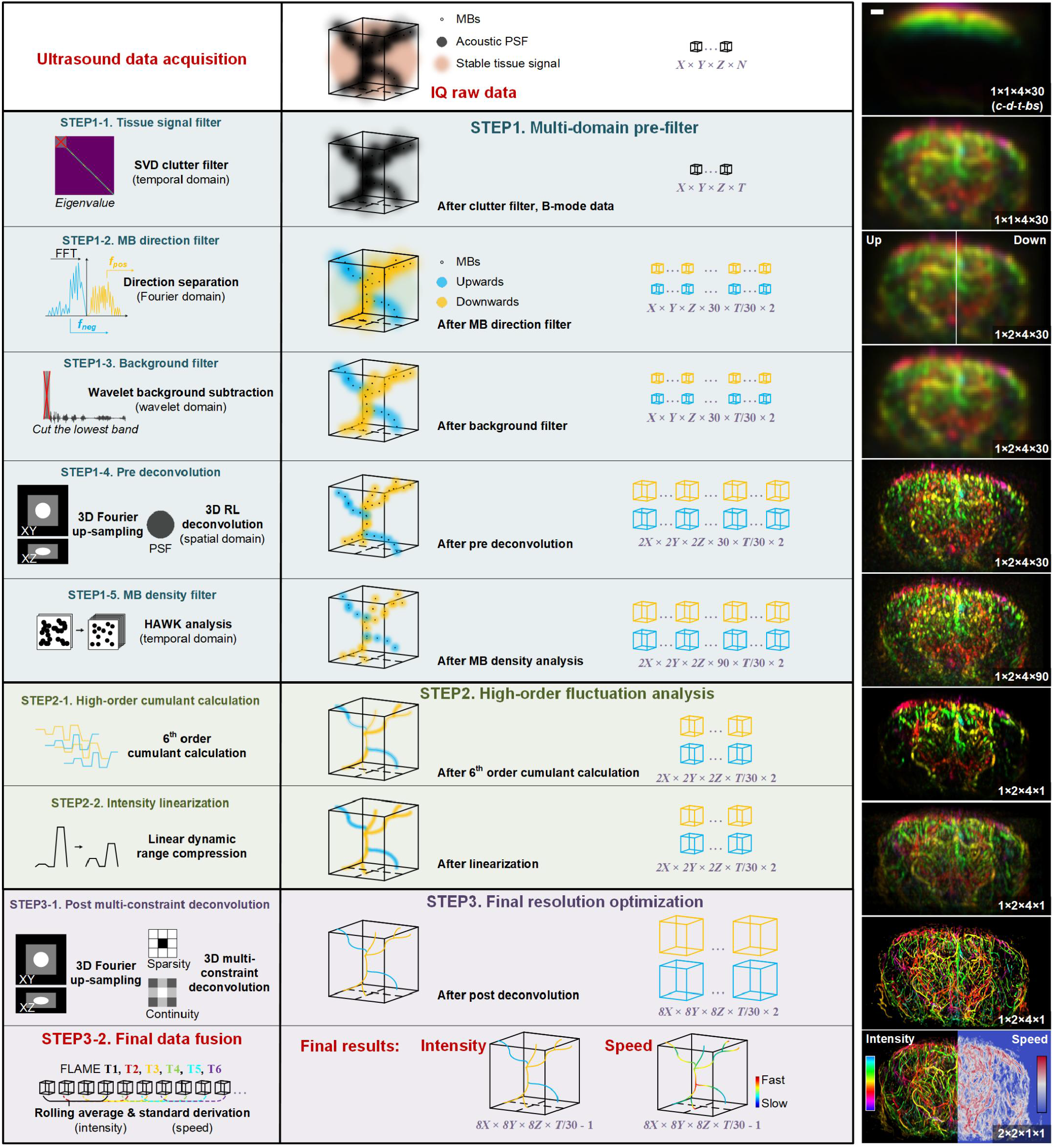
Detailed workflow of FLAME and intermediate results. Left: Schematic diagrams for each processing step; Middle: Conceptual reconstruction results of each step; X, Y, Z, and N represent raw spatiotemporal pixel numbers of acquired US signal, while T denotes the temporal pixel number after stabilization excluding frames with significant motions. We typically use 30 volumes (T/30) per super-resolution reconstruction and 120 volumes as one final FLAME result. Thus, after rolling average, we obtain ‘T/30 – 1’ FLAME volumes. Right: Intermediate reconstruction results of each step. *c*-*d*-*t-bs* represents the number of intensity/speed channels, flow directions, time points, and batch size. Scale bars, 1 mm.

**Extended Data Fig. 2.**
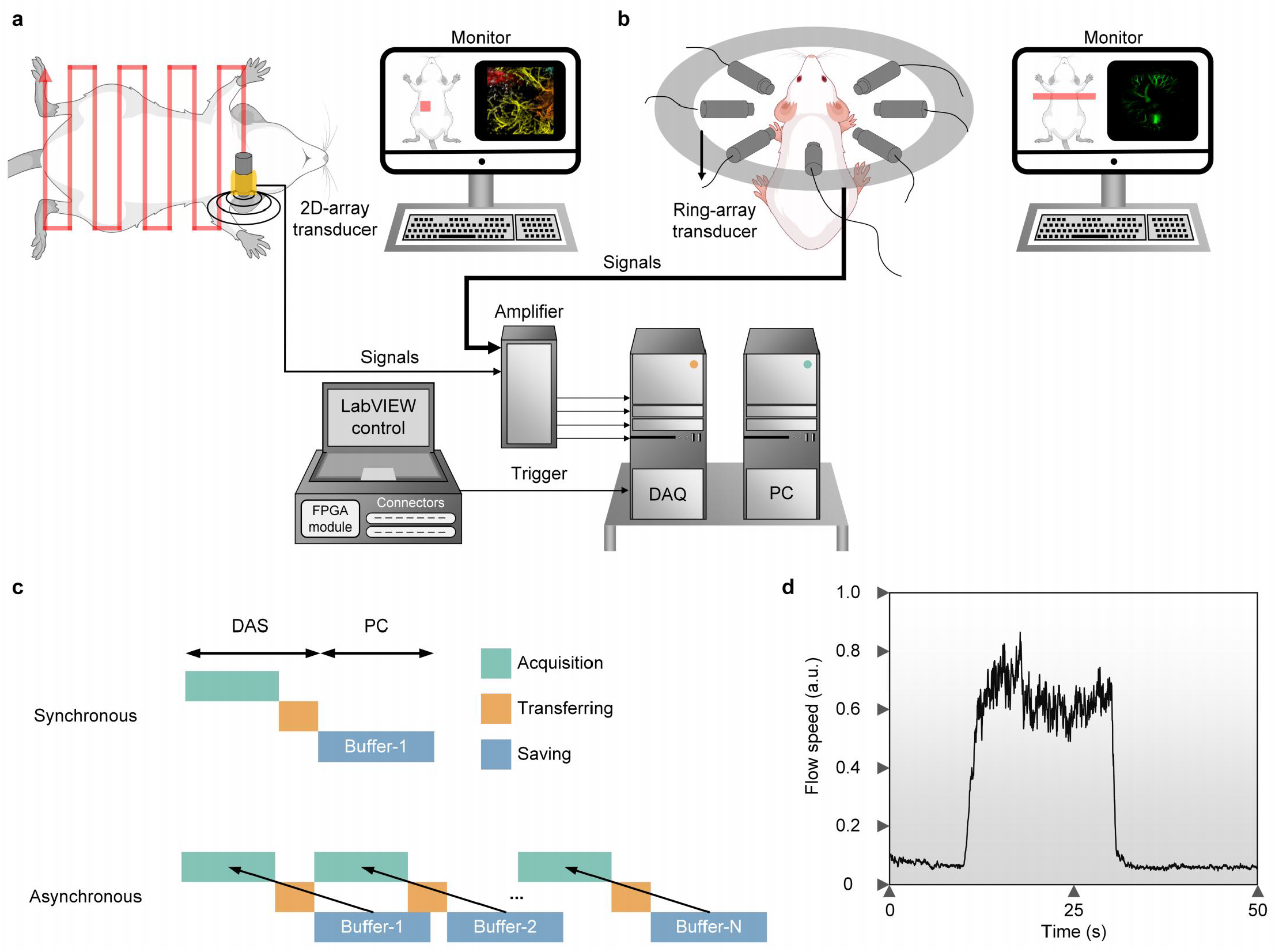
CEUS imaging setups. (**a, b**) Schematic diagrams of two volumetric US imaging setups: the primary setup used throughout this work (**a**), and an alternative setup used exclusively in Fig. 5g–5j and Extended Data Fig. 10 (**b**). **c**, Illustration of the “ping-pong” saving scheme for high-throughput US acquisition at 1.2 kHz. **d**, FLAME at ∼40 Hz visualizing flow dynamics in a MB-perfused silicone tube phantom.

**Extended Data Fig. 3.**
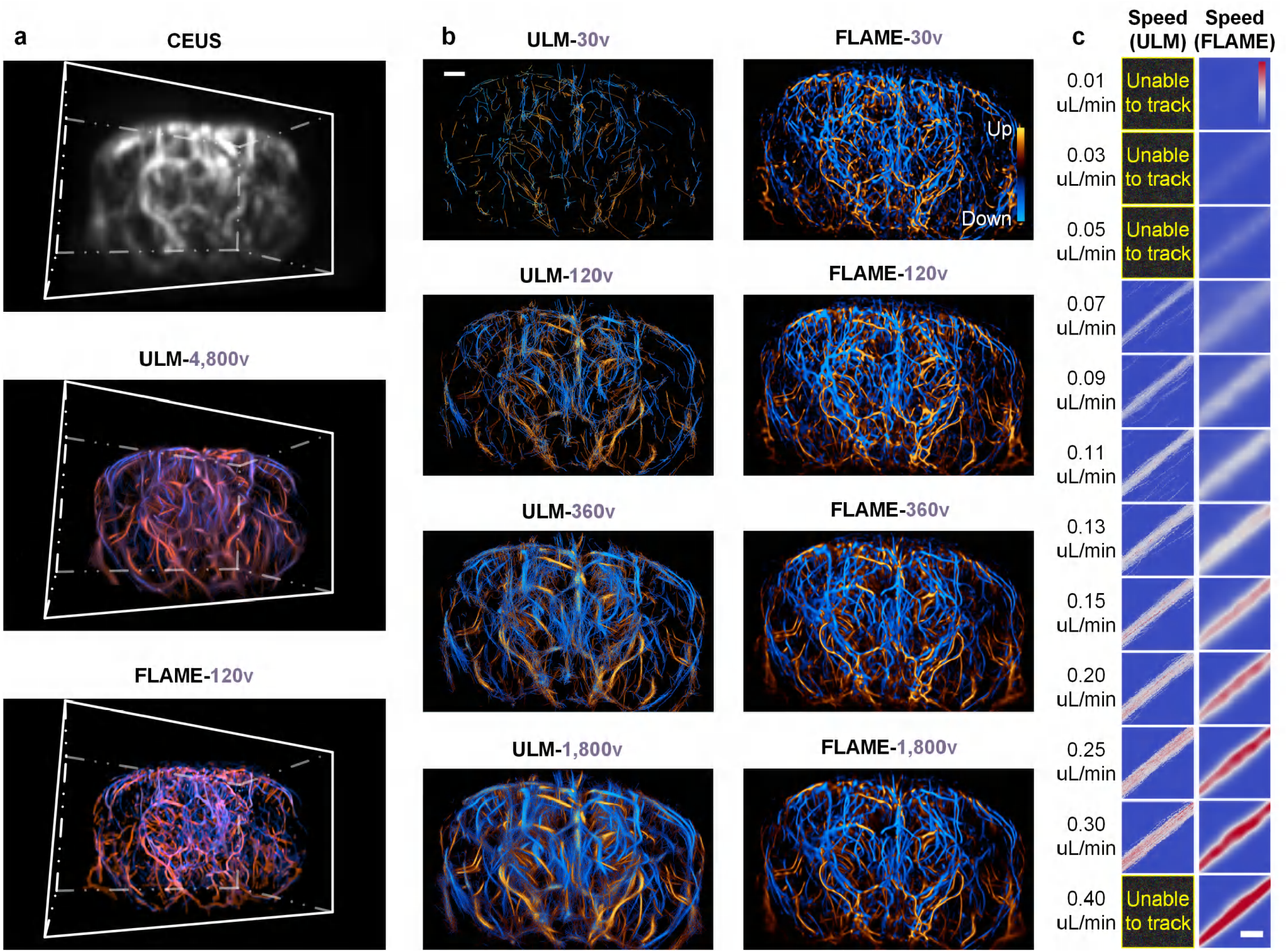
Full visualization of data from Fig. 2. **a**, 3D renderings of volumes reconstructed by CEUS (top), ULM-4,800v (middle), and FLAME-120v (bottom) (*c.f*., Fig. 2a). **b**, Dual-color-coded maximum intensity projection (MIP) views of ULM (left) and FLAME (right) reconstructions using gradually increased numbers of volumes, from top to bottom (*c.f*., Fig. 2a). Scale bars, 1 mm. **c**, Full data representation of Fig. 2f. Scale bars: 0.5 mm.

**Extended Data Fig. 4.**
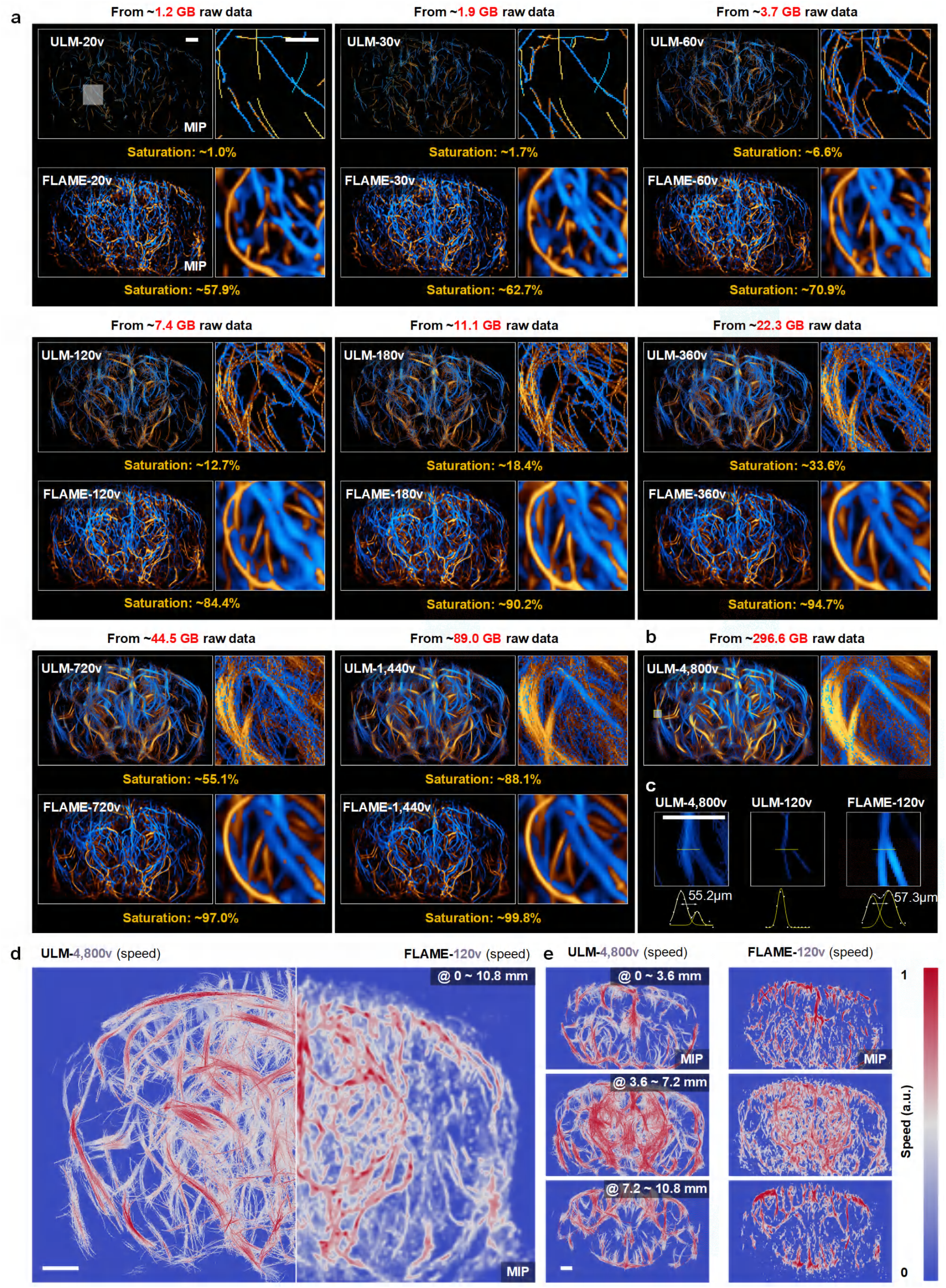
Systematic comparisons of ULM and FLAME reconstructed by different numbers of volumes (*c.f*., Fig. 2a). (**a**) ULM (top) and FLAME (bottom) results reconstructed by different numbers of volumes (20, 30, 60, 120, 180, 360, 720, and 1,440 volumes). Saturation rates and reconstruction-used raw data size were labeled at the bottom and top, respectively. Scale bars, 1 mm (left) and 0.5 mm (right). **b**, ULM result reconstructed by 4,800 volumes. **c**, Zoomed views (top) of ULM-4,800v, ULM-120v, and FLAME-120v from the yellow boxed region in (**b**) and their intensity profiles and multiple Gaussian fitting (bottom) for the structures of microvessels indicated by the yellow lines. The numbers indicate the distance between peaks. Scale bars, 0.5 mm. **d**, Maximum speed projections of ULM-4,800v (left) and FLAME-120v (right) across 0∼10.8 mm. Scale bars, 1 mm. **e**, Maximum speed projections of ULM-4,800v (left) and FLAME-120v (right) at different axial ranges. Scale bars, 1 mm.

**Extended Data Fig. 5.**
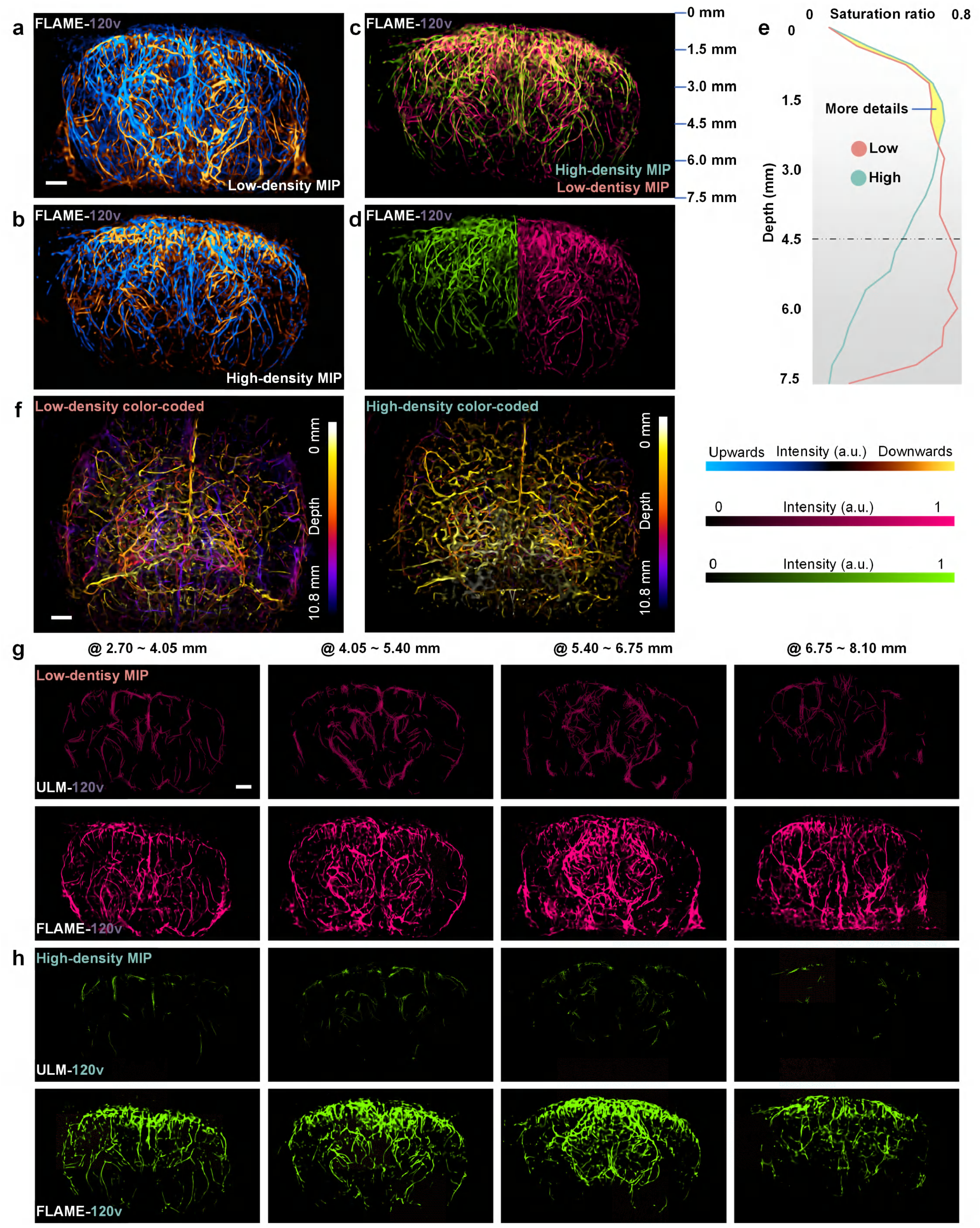
Comparisons of ULM and FLAME in the same mouse under different injected MB concentrations. (**a, b**) Color-coded 3D distributions under FLAME results of mouse-brain microvasculature in Fig. 2b by low MB concentration (**a**) and high MB concentration (**b**). Scale bar, 1 mm. **c**, Merged MIP views of FLAME reconstructions under low MB concentration (salmon) and high MB concentration (turquoise). **d**, MIP views of FLAME reconstructions under high MB concentration (turquoise, left) and low MB concentration (salmon, right). **e**, Saturation ratios of FLAME reconstructions at different z-axis positions under different MB concentrations. **f**, Color-coded 3D distributions of FLAME reconstructions under low MB concentration (left) and high MB concentration (right). Scale bar, 1 mm. (**g, h**) MIP views of ULM-120v (top) and FLAME-120v (bottom) across different z-axis ranges under low MB concentration (**g**) and high MB concentration (**h**). Scale bar, 1 mm.

**Extended Data Fig. 6.**
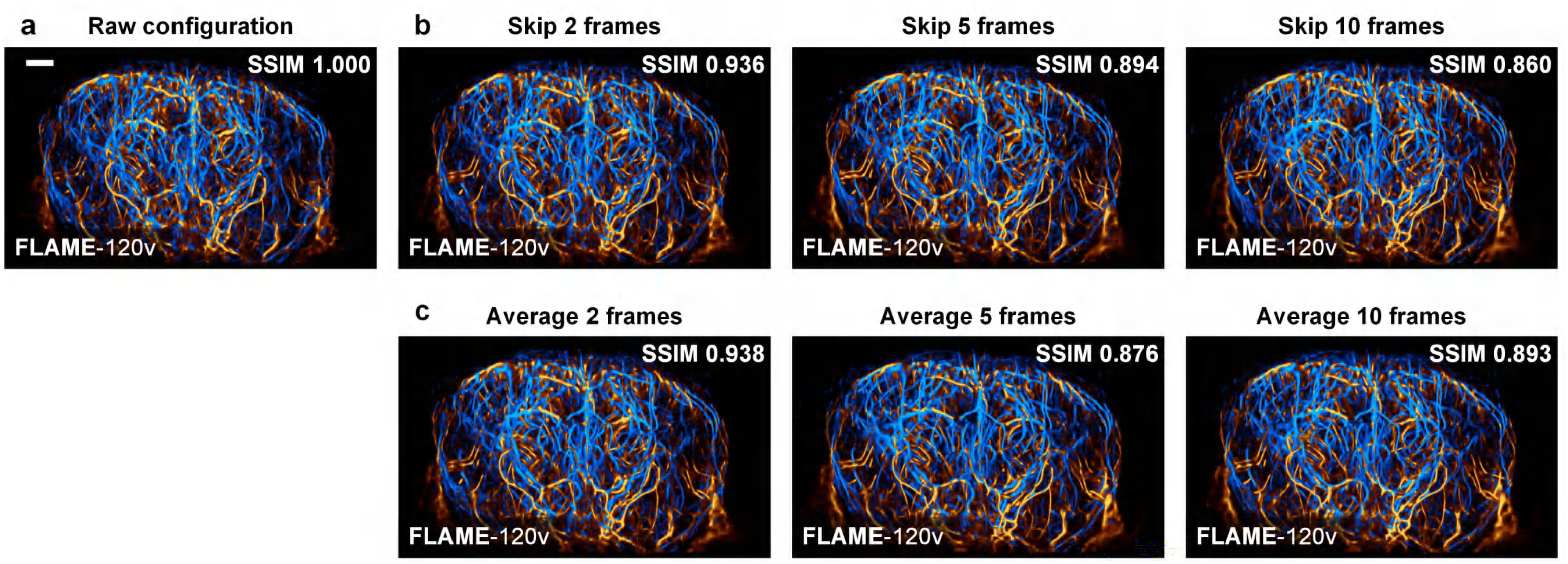
FLAME reconstructions under different experimental configurations. **a**, FLAME-120v under original configuration (identical to **Fig. 2a**). Scale bars, 1 mm. **b**, FLAME-120v under different numbers of artificially skipped volumes (simulating different imaging rates). **c**, FLAME-120v under different numbers of artificially averaged volumes (miming different MB concentrations). SSIM values are labeled on the bottom right corners.

**Extended Data Fig. 7.**
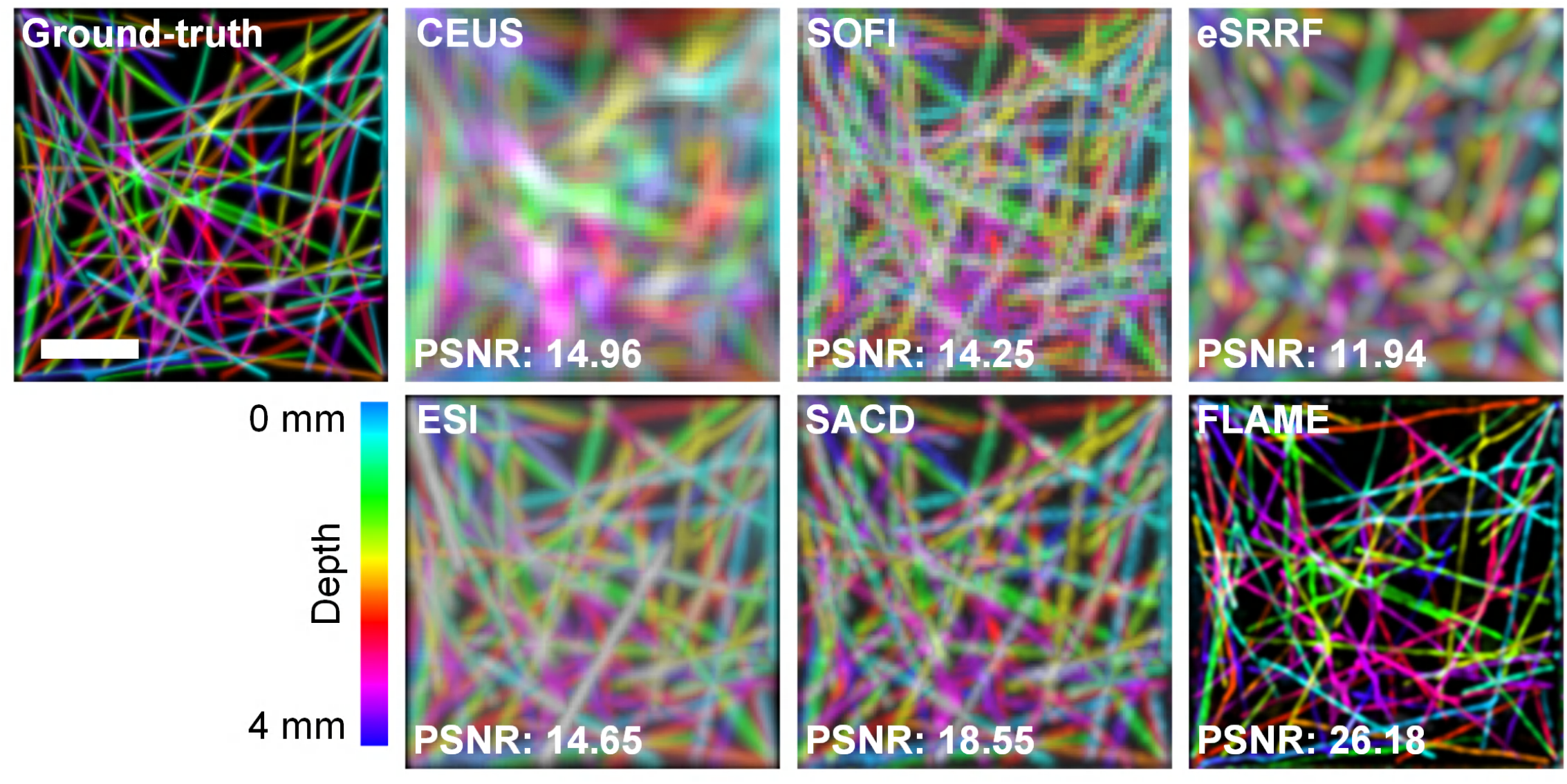
Simulation comparisons of FLAME against different fluorescence fluctuation-based super-resolution methods. SOFI: super-resolution fluctuation imaging^9^; eSRRF: enhanced super-resolution radial fluctuations^24^; ESI: entropy-based super-resolution imaging^23^; SACD: super-resolution imaging based on autocorrelation with two-step deconvolution^11^. Notably, all reconstructions were executed on the SVD-filtered data with 120 volumes. Scale bars: 1 mm.

**Extended Data Fig. 8.**
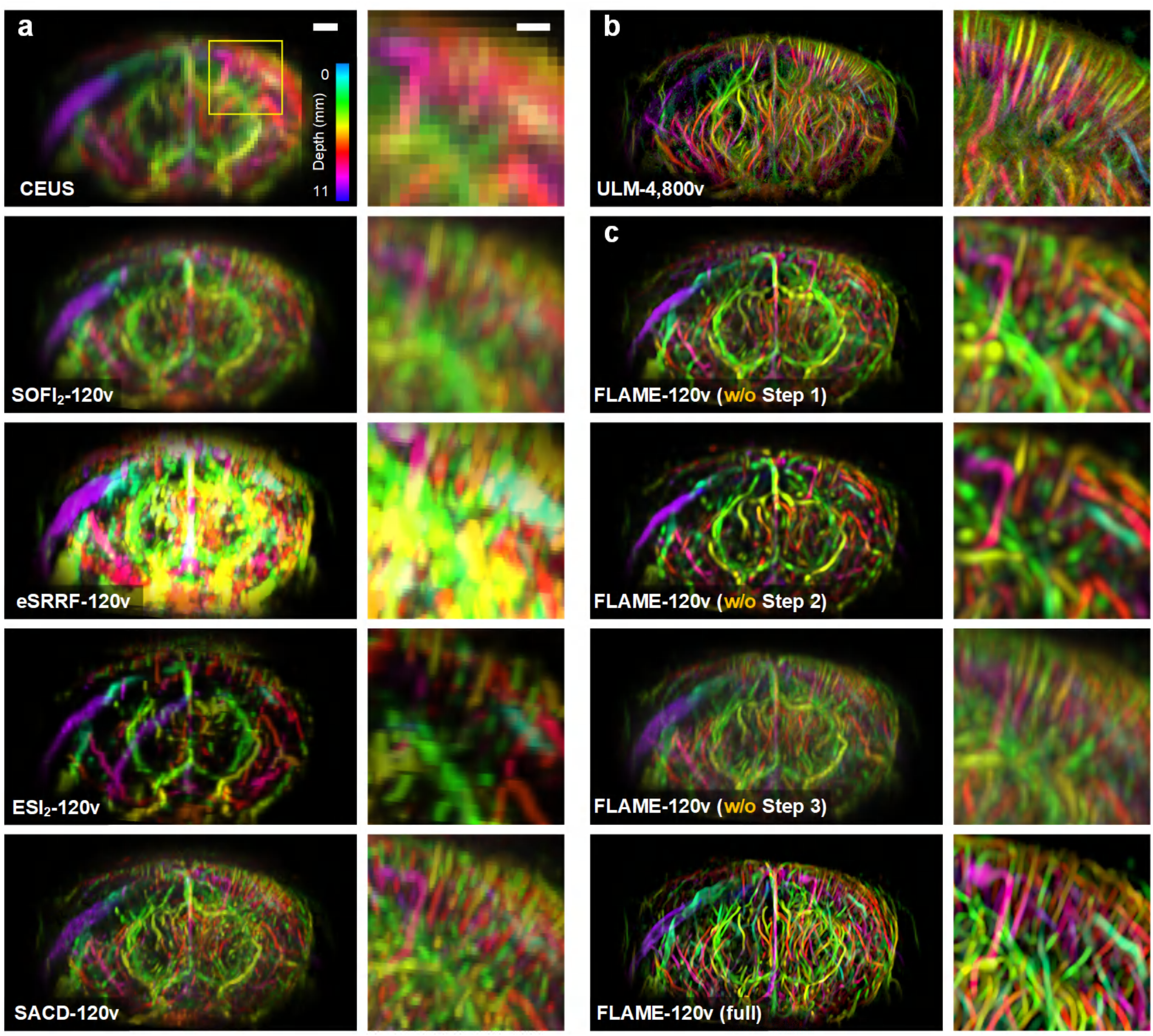
Experimental comparisons of FLAME against different fluorescence fluctuation-based super-resolution methods (*c.f*., Fig. 3a). **a**, From top to bottom: Color-coded volumes (left) and zoomed views from the yellow boxed region (right) of CUES, SOFI, eSRRF, ESI, and SACD. Scale bars: 1 mm (left) and 0.5 mm (right). **b**, ULM color-coded result accumulated by 4,8000 volumes. **c**, FLAME results by removing individual major steps. To avoid canceling the resolution improvement of the cumulant, the ‘FLAME-120v (w/o Step 3)’ image was directly visualized by brightness adjustment. Notably, all reconstructions were executed on the SVD-filtered data with 120 volumes.

**Extended Data Fig. 9.**
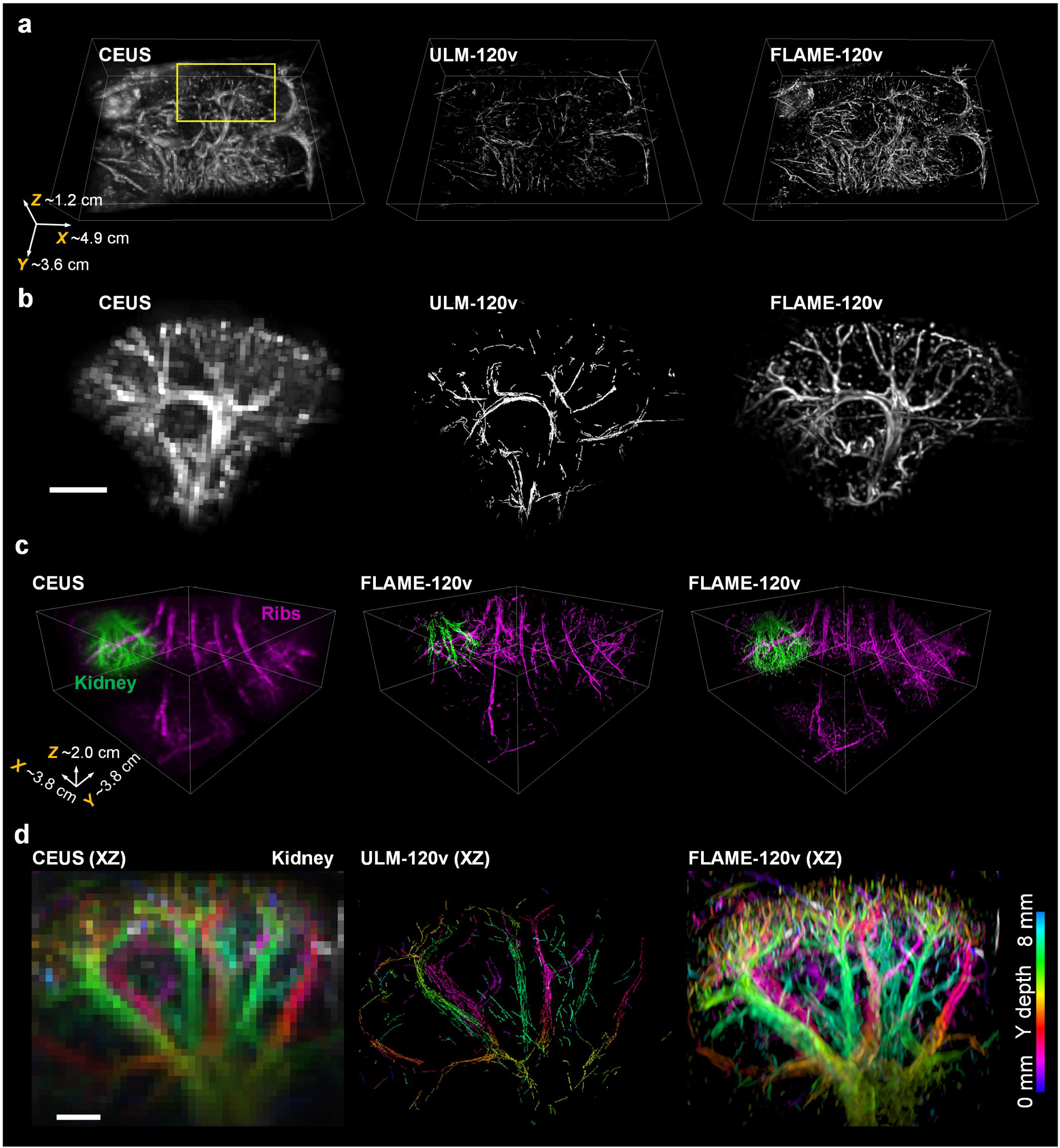
Full data visualization of microvascular system in mice and rats (*c.f*., Fig. 5a-5f). **a**, 3D rendering of whole-body microvasculature in a living mouse recorded by CEUS (left), ULM-120v (middle), and FLAME-120v (right). Volume size is labeled at the bottom left corner. **b**, Magnified 3D rendering results from the yellow box in (**a**). Scale bar, 2 mm. **c**, 3D rendering of renal-region microvasculature in a living rat recorded by CEUS (left), ULM-120v (middle), and FLAME-120v (right). Volume size is labeled at the bottom left corner. **d**, Zoomed color-coded projection views from the green color (kidney) region in (**c**). Scale bar, 2 mm.

**Extended Data Fig. 10.**
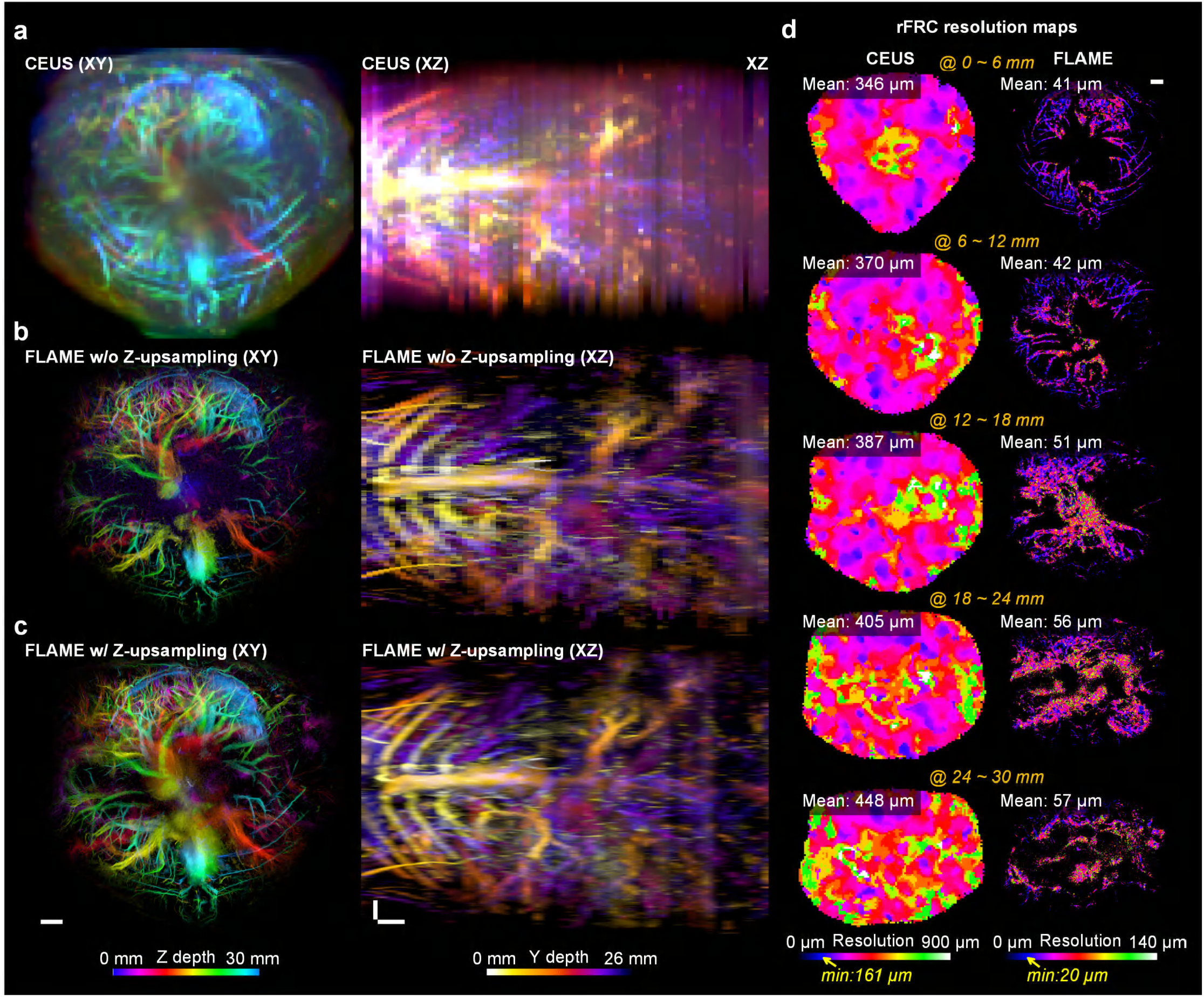
Full visualization and resolution evaluation of data from Fig. 5h-5j. **a**, Color-coded horizontal (left, XY projected) and vertical (right, XZ projected) views of the primary torso microvasculature in a living mouse recorded by CEUS. (**b, c**) FLAME results from data in (**a**) without (**b**) and with (**c**) upsampling along the z axis. As shown in Extended Data Fig. 2b, transverse imaging across the mouse torso was performed using a ring-array US transducer to capture cross-sectional structures. For recording the 3D vascular distribution, the imaging stage was translated along the torso, acquiring 2D data at each position. To accelerate volumetric imaging and minimize motion blur, the scanning step size along the z-axis was set to 0.4 mm. To compensate for the reduced spatial resolution, we applied an additional ∼4× Fourier upsampling along the z-axis during post-processing. Scale bars, 2 mm. **d**, rFRC resolution maps of CEUS and FLAME results at different z-axis positions (from top to bottom). Mean resolutions are labeled at the top left corners. Scale bars, 2 mm.

