## Supplementary Information for "High-throughput 3D super-resolution ultrasound imaging"

### Supplementary Figures.

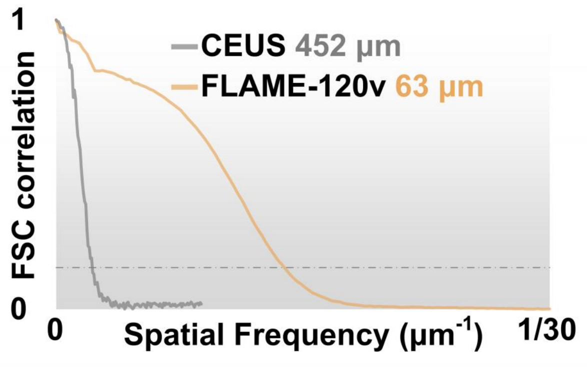


**Supplementary Fig. 1 | FSC resolution evaluation (*c.f.*, Fig. 3a).**

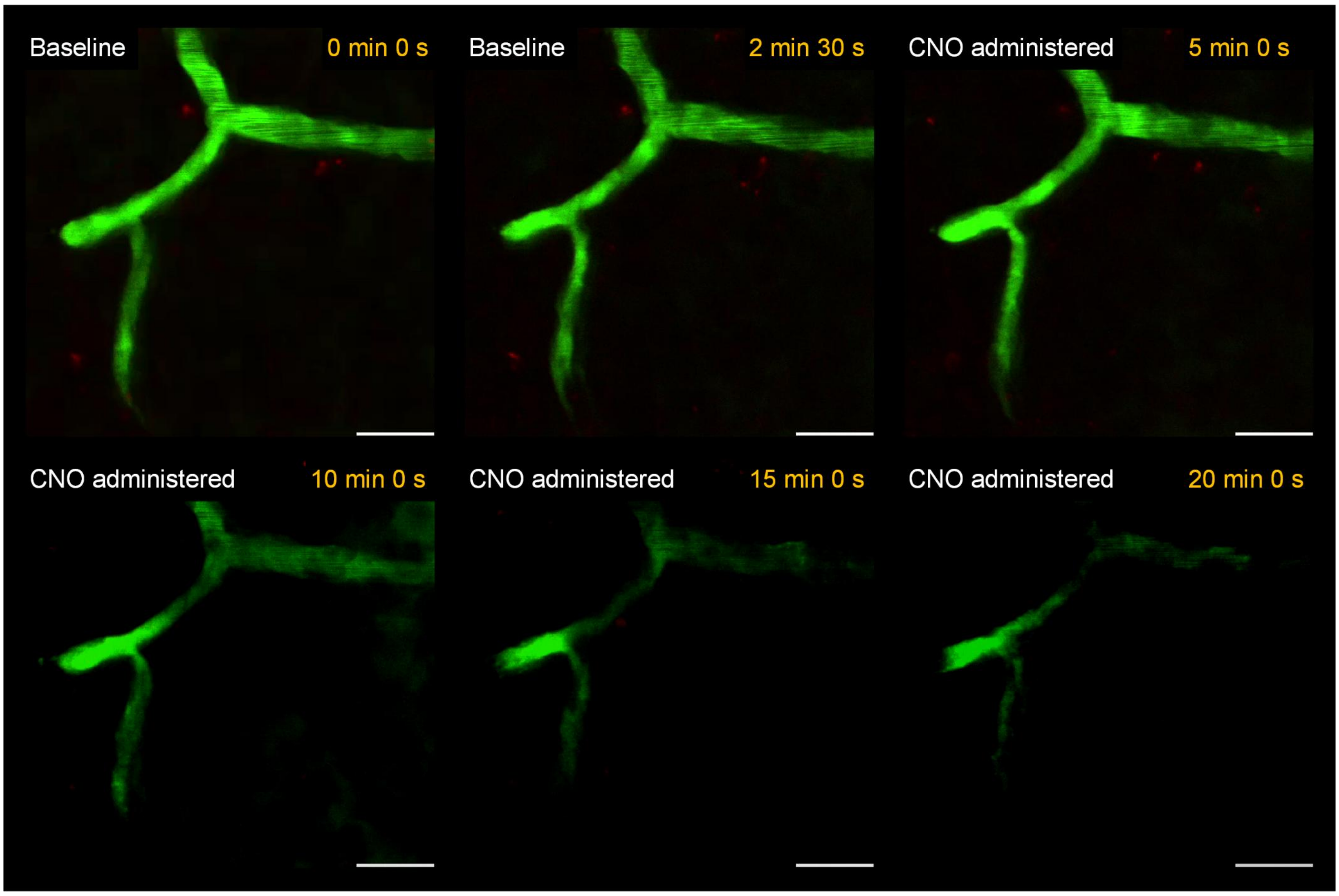


**Supplementary Fig. 2 | Representative two-photon fluorescence microscopy snapshots cross-validate the effects of CNO on cortical microvessels in a transgenic mouse model.** Scale bars, 20 µm.

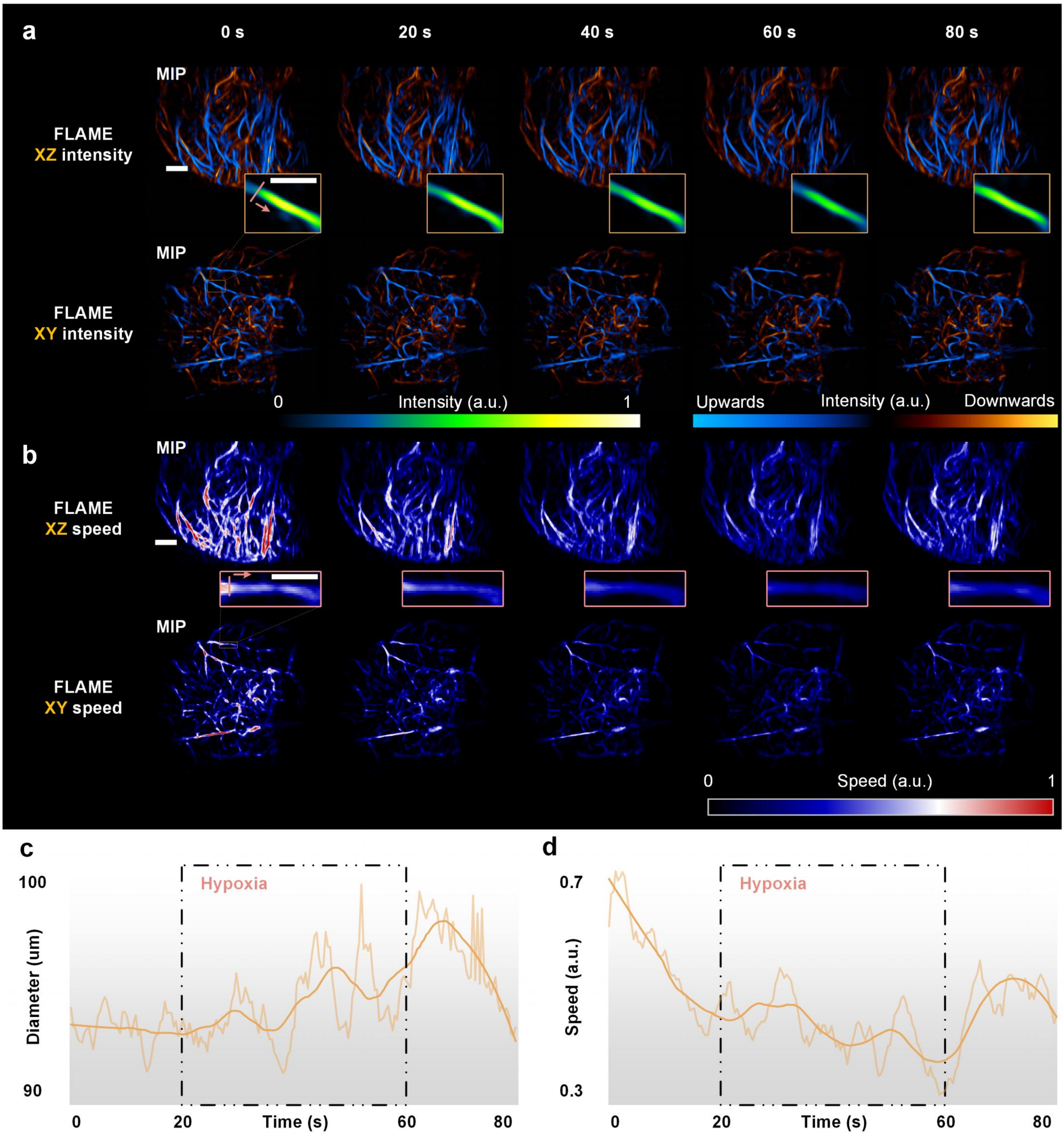


**Supplementary Fig. 3 | FLAME captures the hypoxia responses in the cerebral microvasculature of a living mouse with intact scalp and skull. a**, Representative maximum intensity projections at 5 time points under XZ (top) and XY (bottom) views. Scale bars: 1 mm (top) and 0.5 mm (bottom). **b**, Representative maximum flow speed projections at 5 time points under XZ (top) and XY (bottom) views. Insets showcase an individual vessel from the yellow-boxed region at 5 different time points. Scale bars: 1 mm (top) and 0.5 mm (bottom). (**c**, **d**) Average diameter (**c**) and speed (**d**) of the selected vessel over time. Translucence lines are smoothed data after averaging. The hypoxia process induced vessel diameter and speed reactions (diameter expansion followed by contraction; speed decrease followed by increase) were successfully detected by our FLAME.

### Captions for Supplementary Videos.

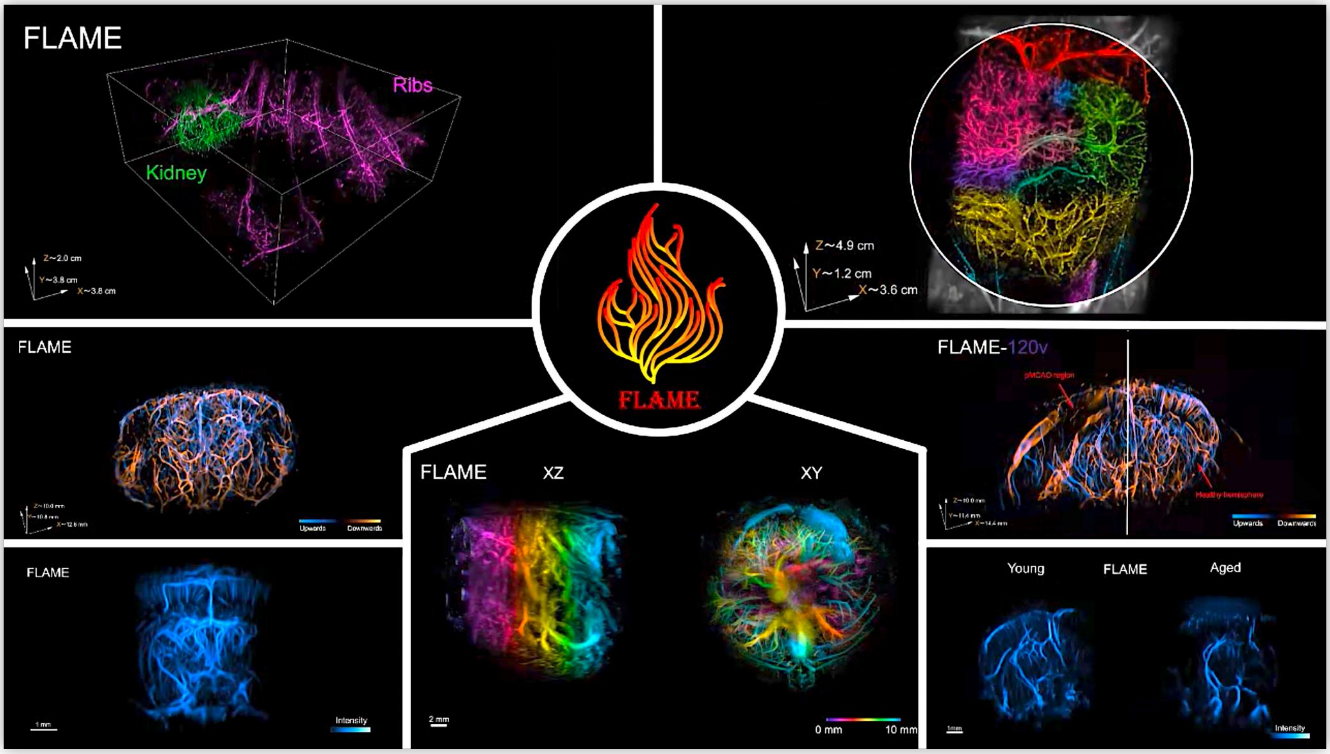


**Supplementary Video 1 | FLAME overview.** Video abstract illustrates the main concepts behind FLAME: (1) the experimental operation based on conventional CEUS imaging; (2) the reconstruction workflow; (3) the conceptual differences between ULM and FLAME; (4) the representative imaging results.

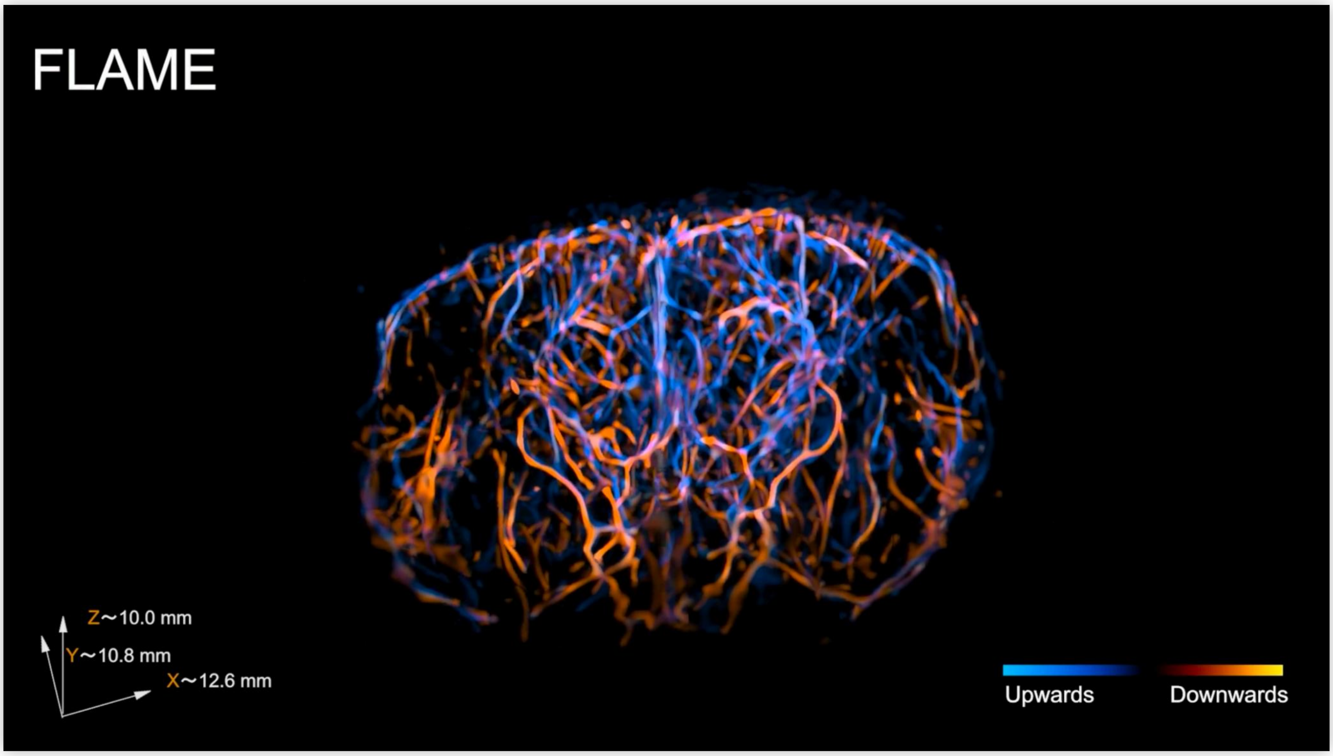


**Supplementary Video 2 | FLAME resolves brain-wide microvasculature.** Whole-brain microvascular network in a mouse with intact scalp and skull (*c.f.*, **Fig. 2a**). Part I the shows the 3D rendering views. Part II presents the color-coded projection views.

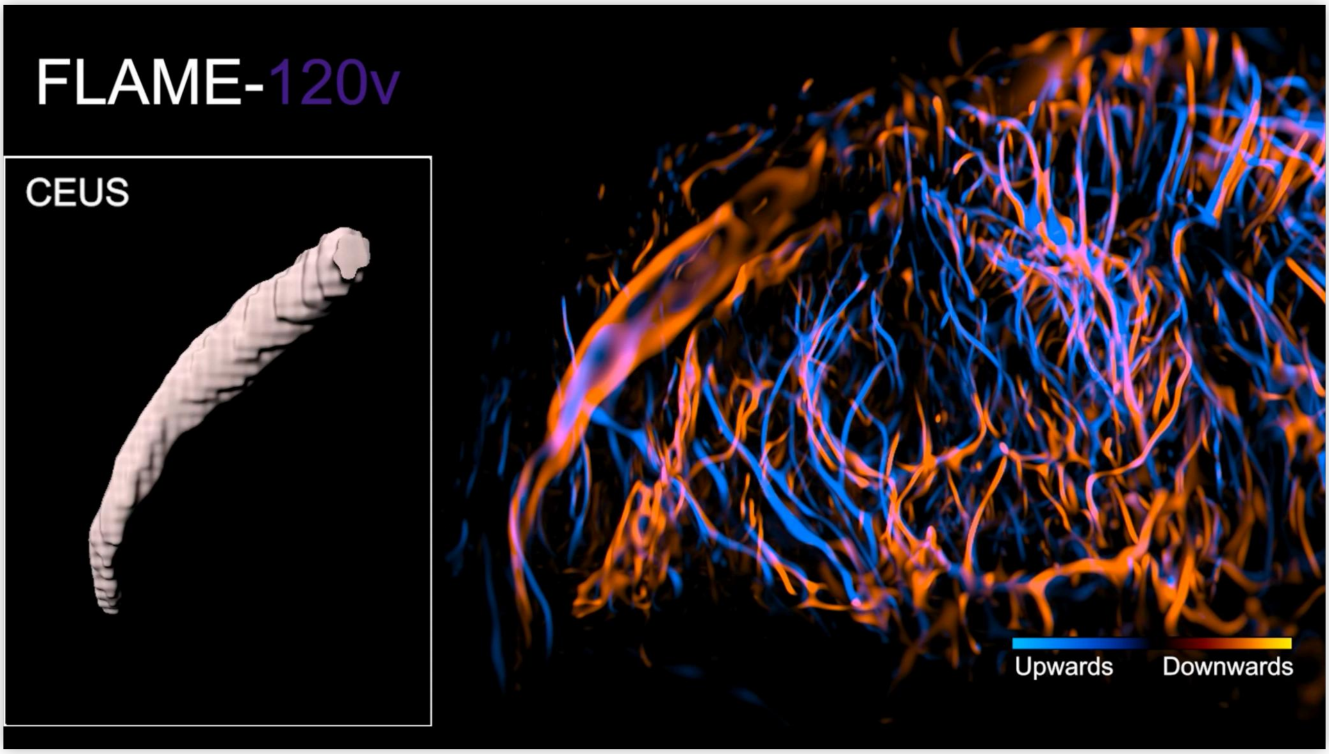


**Supplementary Video 3 | FLAME characterizes stroke in mice.** Whole-brain microvascular network in a pMCAO model mouse (*c.f.*, **Fig. 3a**). The 3D rendering shows the differences between healthy and stroke hemispheres. By isosurface-rendering, FLAME clearly resolves three intersected large vessels with well-defined boundaries of topological structures, where both CEUS and ULM results appear blurry and less discernible.

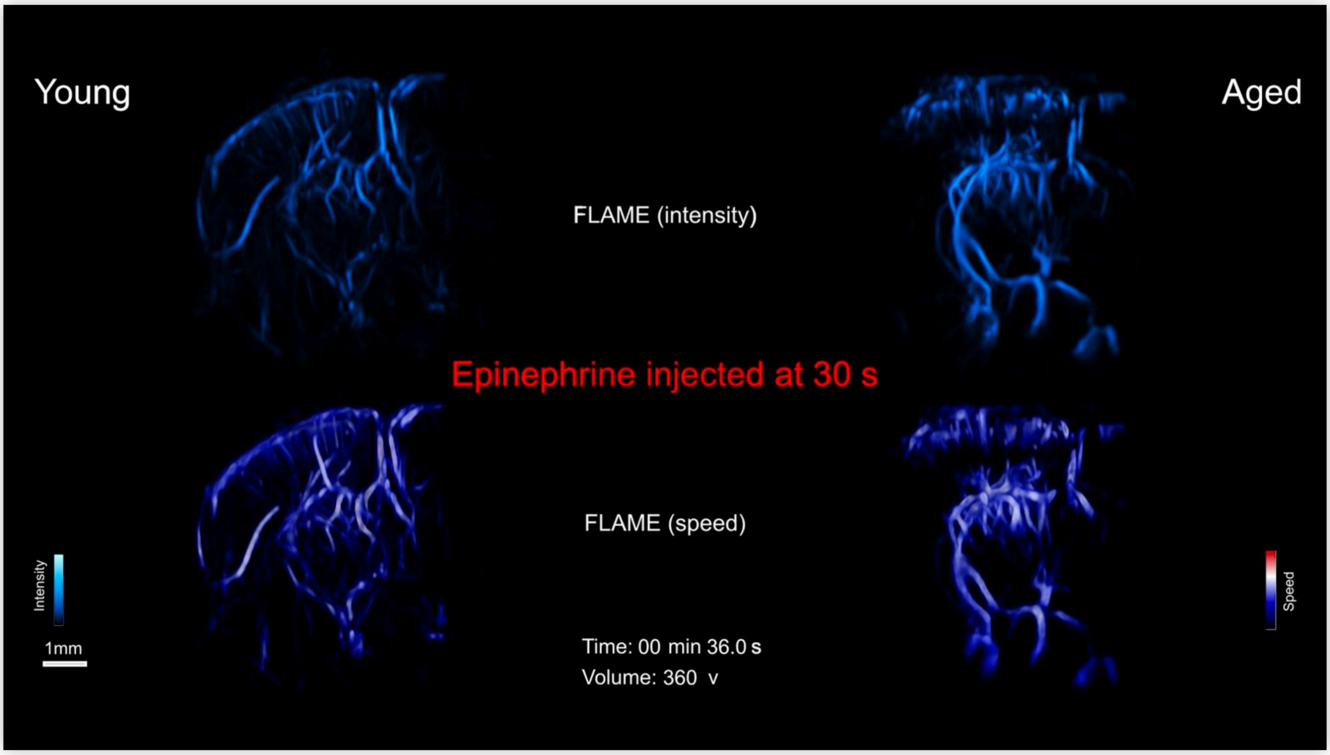


**Supplementary Video 4 | FLAME monitors acute hemodynamic reactions of mouse brains with different ages.** Differential regression patterns in perfusion and flow speed between young and aged mouse brains (*c.f.*, **Fig. 4c**). Part I shows the 3D renderings of microvascular network imaged by CEUS and FLAME. Part II illustrates the temporal evolution of blood perfusion (top) and flow speed (bottom) in young and aged mouse brains, accompanied by magnified views of the cortical regions. Young mice respond quickly and transiently, while aged mice show a sustained response without recovery after epinephrine injection at 30 s.

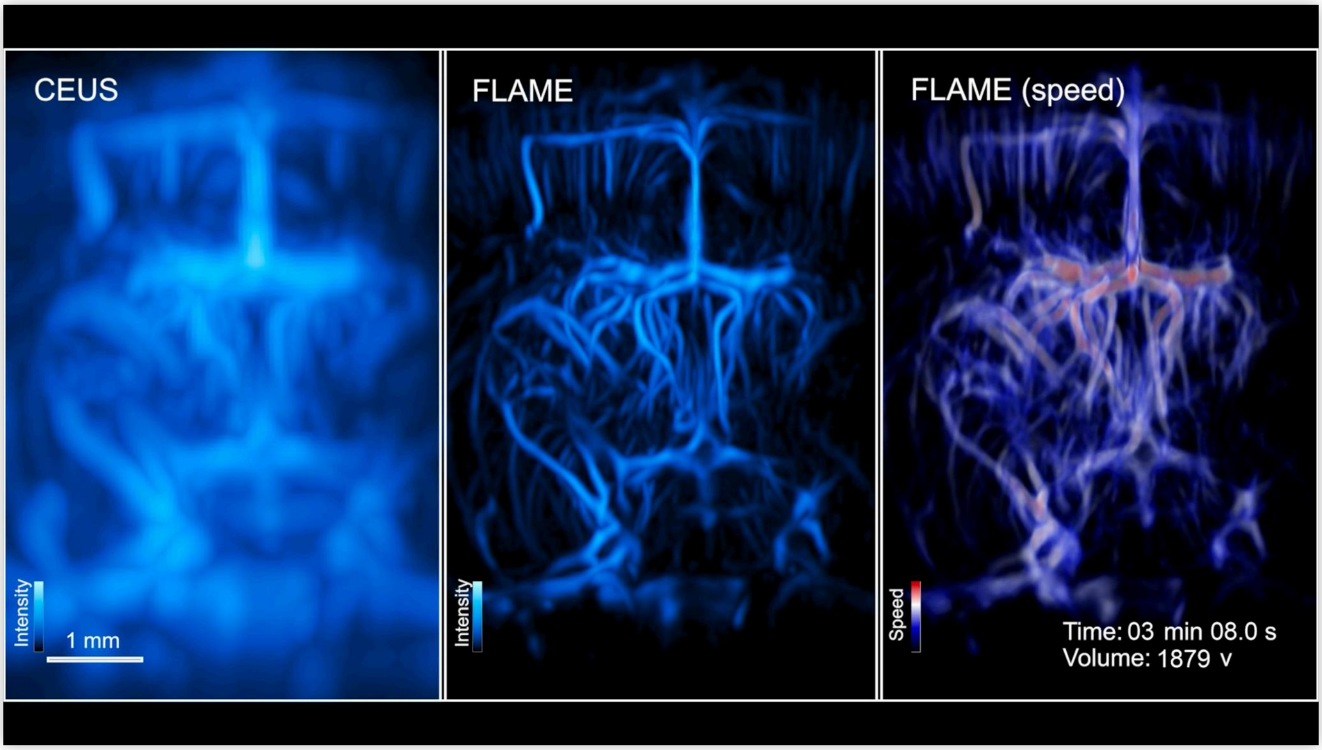


**Supplementary Video 5 | FLAME reveals vascular changes by chemogenetic stimulation.** Local vascular changes by chemogenetic stimulation (*c.f.*, **Fig. 4i**). Part I shows the 3D rendering views of microvascular network before injection under CEUS and FLAME. Part II illustrates the temporal evolution of blood perfusion (top) and flow speed (bottom), accompanied by magnified views of the cortical regions. FLAME reveals a localized decrease in capillary perfusion, whereas major vessels exhibit largely sustained perfusion following CNO injection at 60 s.

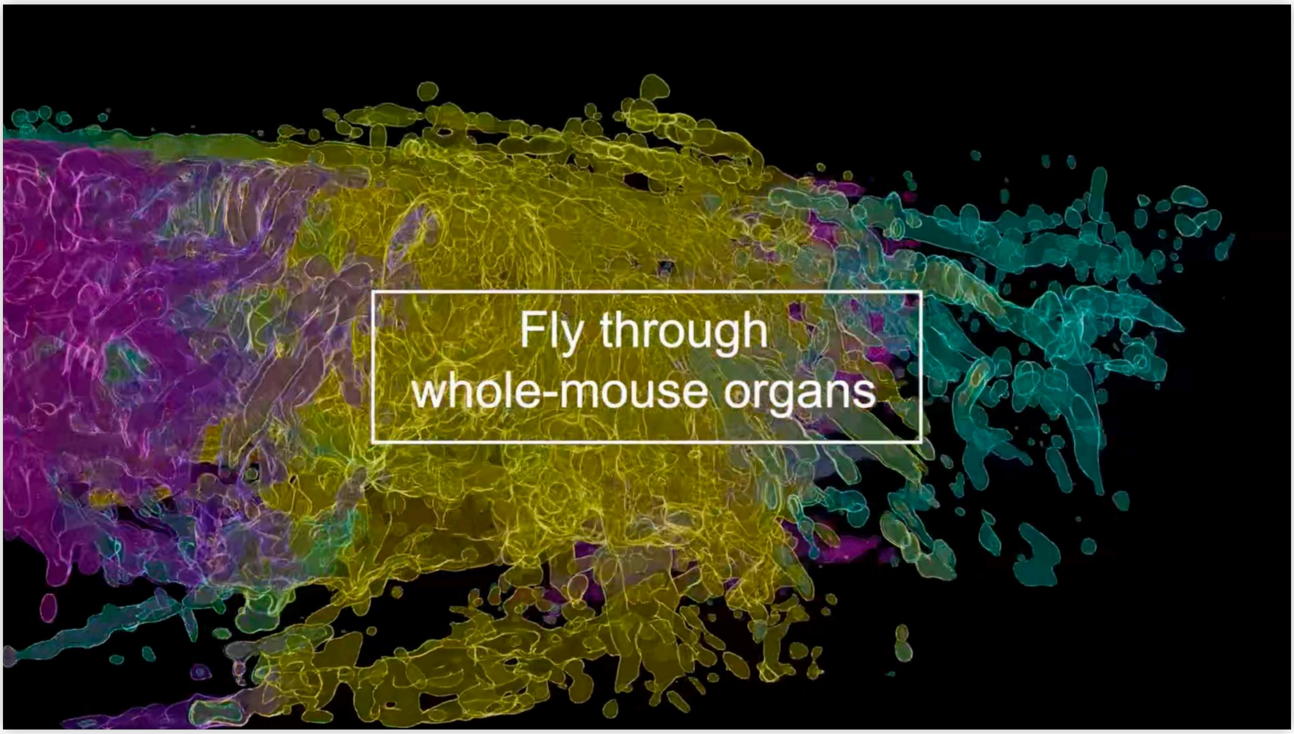


**Supplementary Video 6 | FLAME records whole-body microvascular system.** High-throughput imaging of whole mouse body including heart, bladder, liver, stomach, spleen, intestine, etc. (*c.f.*, **Fig. 5b**). Part I shows the 3D rendering comparison of CEUS and FLAME. Part II presents a fly-through visualization of organ dissection in the mouse.

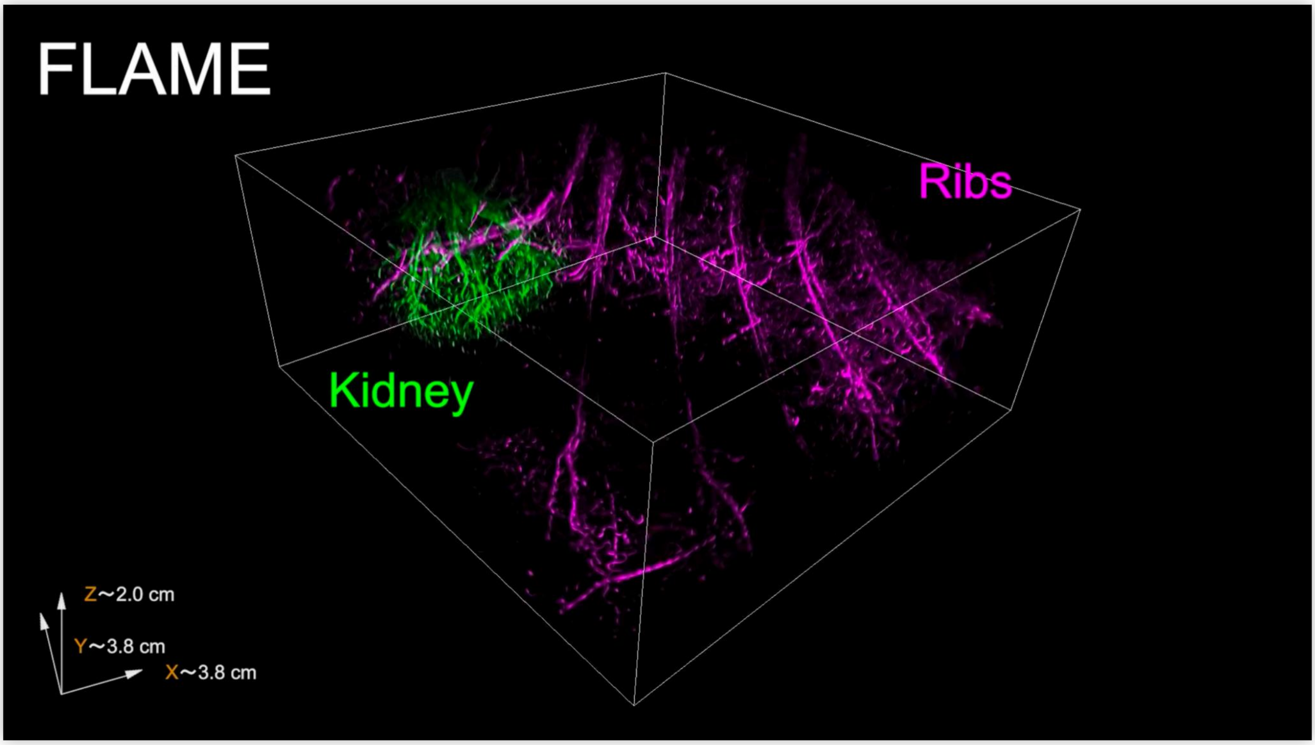


**Supplementary Video 7 | FLAME tracks vasculature in deep rat organs beyond 2 cm depth.** Renal region microvasculature in a living rat (*c.f.*, **Fig. 5e**).

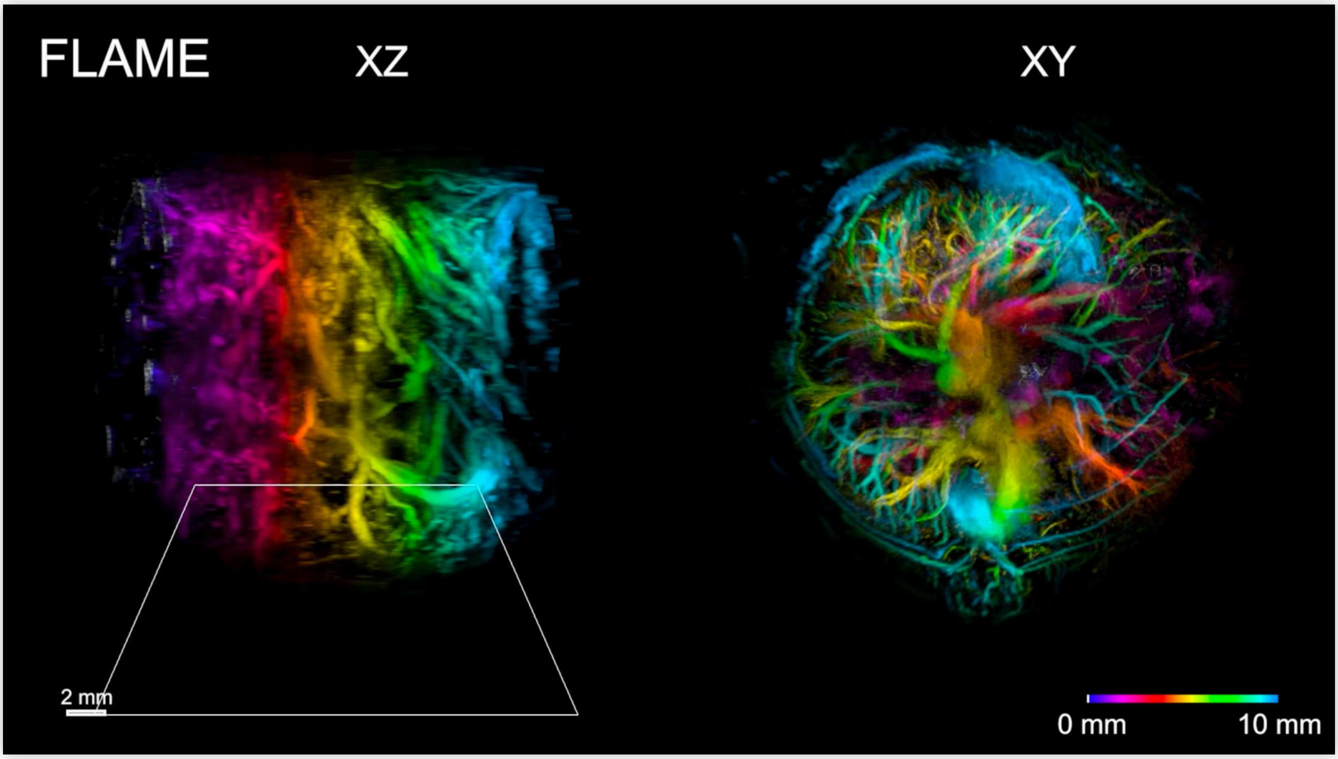


**Supplementary Video 8 | FLAME visualizes the primary torso microvasculature.** Cross-sectional imaging along the torso in a living mouse (*c.f.*, **Fig. 5j**) under CEUS and FLAME.
